# Condition-adaptive fused graphical lasso (CFGL): an adaptive procedure for inferring condition-specific gene co-expression network

**DOI:** 10.1101/290346

**Authors:** Yafei Lyu, Lingzhou Xue, Feipeng Zhang, Hillary Koch, Laura Saba, Katerina Kechris, Qunhua Li

## Abstract

Co-expression network analysis provides useful information for studying gene regulation in biological processes. Examining condition-specific patterns of co-expression can provide insights into the underlying cellular processes activated in a particular condition. One challenge in this type of analysis is that the sample sizes in each condition are usually small, making the statistical inference of co-expression patterns highly underpowered. A joint network construction that borrows information from related structures across conditions has the potential to improve the power of the analysis.

One possible approach to constructing the co-expression network is to use the Gaussian graphical model. Though several methods are available for joint estimation of multiple graphical models, they do not fully account for the heterogeneity between samples and between co-expression patterns introduced by condition specificity. Here we develop the condition-adaptive fused graphical lasso (CFGL), a data-driven approach to incorporate condition specificity in the estimation of co-expression networks. We show that this method improves the accuracy with which networks are learned. The application of this method on a rat multi-tissue dataset and The Cancer Genome Atlas (TCGA) breast cancer dataset provides interesting biological insights. In both analyses, we identify numerous modules enriched for Gene Ontology functions and observe that the modules that are upregulated in a particular condition are often involved in condition-specific activities. Interestingly, we observe that the genes strongly associated with survival time in the TCGA dataset are less likely to be network hubs, suggesting that genes associated with cancer progression are likely to govern specific functions, rather than regulating a large number of biological processes. Additionally, we observed that the tumor-specific hub genes tend to have few shared edges with normal tissue, revealing tumor-specific regulatory mechanism.

**Author summary:** Gene co-expression networks provide insights into the mechanism of cellular activity and gene regulation. Condition-specific mechanisms may be identified by constructing and comparing co-expression networks of multiple conditions. We propose a novel statistical method to jointly construct co-expression networks for gene expression profiles from multiple conditions. By using a data-driven approach to capture condition-specific co-expression patterns, this method is effective in identifying both co-expression patterns that are specific to a condition and that are common across conditions. The application of this method on real datasets reveals interesting biological insights.

## Introduction

Gene co-expression network analysis is a useful tool for studying the complex regulatory machinery in organisms [1][2][3][4]. When the gene expression profiles under multiple conditions are available, comparing co-expression networks across conditions could reveal co-expression patterns that are common across conditions and those that are unique to a condition [5][6][7][8][9], providing insights on how genes work together to regulate biological processes under different conditions. It has been demonstrated that complex diseases are likely to be regulated by condition-specific mechanisms while condition-specific hub genes are likely to be drug targets [10][11][12].

The Gaussian graphical model and its variants have been widely used for studying biological networks [13][14][15][16][17][18][19]. This method models the joint distribution of a set of variables and characterizes the conditional dependence between each pair of variables given all the other variables through the precision matrix (a.k.a. inverse covariance matrix) of the joint distribution [20]. Unlike co-expression models based on marginal correlation, e.g.[21], which do not distinguish the direct and indirect (e.g. through intermediate genes) relationship between genes, the direct relationship between a pair of genes can be inferred from the conditional independence estimated from the Gaussian graphical model. Many algorithms have been proposed to obtain a sparse estimate for the precision matrix, for example, graphical lasso [22] and neighborhood selection [23]. These algorithms make it possible to construct gene co-expression networks using graphical models. A graph generated from this estimate, where genes are represented as nodes and entries in the estimated precision matrix as edges, provides a useful tool for visualizing the relationships between genes and for generating biological hypotheses.

In a multi-condition gene expression study, the co-expression profiles across conditions typically are related, for example, due to shared pathways in different tumor subtypes, or common regulatory mechanisms for housekeeping genes in different tissues. A joint analysis that borrows information across conditions potentially can reveal common structures and increase the power of statistical inference, which is especially useful when the sample sizes are small. Recently, several methods have been proposed to jointly analyze multiple graphical models. Meinshausen et al. [24] incorporated a non-convex hierarchical group lasso penalty into the graphical lasso to encourage common 0’s (i.e. absence of edges) in the precision matrix across conditions. Danaher et al. [6] proposed a joint graphical lasso model by adding an additional convex penalty to the graphical lasso objective function. They proposed two choices for the convex penalty: a group penalty that encourages a shared pattern of sparsity and a fused lasso penalty that encourages similarities in both network sparsity and edge weights.

Despite their differences, these methods encourage similarities equally across all edges and all conditions. This inherently assumes that the similarity across conditions is similar for all edges and that the precision matrices in all conditions are equally similar to each other. For gene co-expression networks across different conditions, however, both assumptions are violated due to the heterogeneity across genes and across conditions. First, edges in the networks often have different levels of conservation across conditions. For example, in a network consisting of multiple pathways, the pathways involving basic cellular functions tend to be more conserved across tissues than those involving tissue-specific functions. Second, when there are multiple conditions, some conditions may be more similar to each other than others. For example, tissues with the same embryonic origin may have more similar pathways than those with different origins. More recently, several methods have been proposed to allow more structural heterogeneity in joint estimation. Zhu et al. [25] introduced a non-convex truncated *l*_1_ penalty on the pairwise differences between the precision matrices to encourage elementwise clustering of similar entries across conditions. To incorporate external information on shared subgraphs across conditions, Ma et al. [26] grouped edges shared across conditions based on external information and extended the neighborhood selection method to a joint analysis with the proposed penalty. To handle heterogeneity in similarities across conditions, Seagusa et al. [27] proposed a Laplacian shrinkage penalty to incorporate the pairwise distance between conditions, and proposed using hierarchical clustering to obtain the pairwise distance when it is unknown a priori. While these methods improve the flexibility in estimation, they do not completely address the issues in studying condition-specific co-expression networks. For example, though the approach in Zhu et al. [25] allows abrupt elementwise difference across conditions, it still implicitly assumes that the majority of edges are common across conditions and penalizes condition-specificity. The approach in Ma et al. [26] relies on the availability and the quality of external information, which is still limited for gene co-expression relationships. The approach in Seagusa et al. [27] uses external information or hierarchical clustering to define the weighted subpopulation network and only partially addressed the issue of condition specificity.

In this work, we proposed an adaptive approach to simultaneously addressing condition specificity and heterogeneity across conditions in the estimation of multiple co-expression networks. Our strategy is to incorporate a binary weight matrix that contains information on whether or not an edge is common between conditions in the Fused Graphical lasso framework. We proposed a strategy to learn this matrix adaptively from the data based on a test for differential co-expression, though it can also be obtained from external sources. The incorporation of this matrix not only accounts for the difference between condition-common edges and condition-specific edges but also makes the estimation adaptive to the distance between different conditions. In this way, one can borrow information across conditions for common edges, while estimating differential edges in a condition-specific manner. We provide a computationally efficient implementation using the alternating direction method of multipliers (ADMM) algorithm. Our simulations show that this method generates more accurate results in both edge detection and edge weight estimation. We applied our method to a rat multi-tissue dataset and a TCGA breast cancer dataset and obtained interesting biological insights.

## Results

### Review of Graphical Lasso model and Fused Graphical Lasso model

We first briefly describe the Graphical Lasso (GL) method [22] and the Fused Graphical Lasso (FGL) method [6]. Suppose the gene expression profiles are available across K conditions, where conditions are, for example, different tissues or disease statuses. Denote the gene expression levels **Y**^(k)^ for the condition k, k.1,2,…,K, as a n_k_ ×p matrix, where p is the number of genes, which is common across all conditions, and n_k_ is the number of observations, which can vary across conditions. Suppose that gene expression levels within each condition, 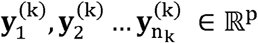, are identically drawn from N(**µ**_k_,**∑**_k_), where **µ**_k ∈_,ℝ^p^ and **∑**_k_ is a positive definite p×p matrix. Then zero entries in the precision matrix 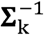 correspond to the pairs of genes that are conditionally independent given all other genes in the dataset. Based on the precision matrix,^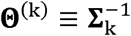^, a gene co-expression network can be constructed by representing the genes as nodes and conditional dependencies as edges in a graph.

The most direct way to analyze such data is to estimate *k*, individual graphical models separately. We can use the graphical lasso method to compute a separate *l*_1_ penalized estimator of 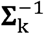 for each condition by solving,

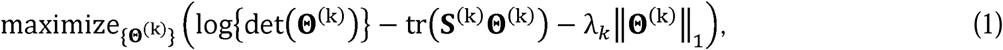

Where **S**^(k)^ = (**y**^(k)^)^T^ **y**^(k)^/n_k_ is the empirical covariance matrixes of **y**^(k)^ and λ_k_‖Θ^(k)^‖_1_ is the penalty terms with λ_k_ as a non-negative tuning parameter and ‖Θ^(k)^‖_1_ as the L_1_ norm of Θ(^k^). However, when the conditions are related, separate estimation ignores the common structure shared across conditions and can also mask differences critical in understanding condition-specificity in the co-expression pattern.

To address this issue, Danaher et al. [6] developed a fused graphical lasso model to jointly estimate multiple graphical models from related conditions. This model incorporates the generalized fused lasso penalty P({Θ}) [28] to the log-likelihood,

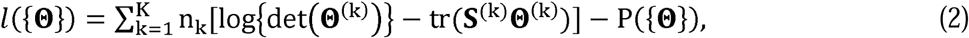

such that information can be borrowed across conditions. The penalty penalty P({Θ}) is a convex penalty with two terms,

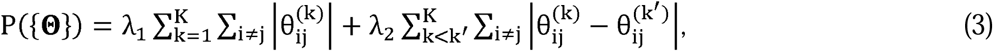

where λ_1_ and λ_2_ are non-negative tuning parameters, and 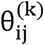 is the (i,j)-th element of the matrix Θ^(k)^. The first term, which is the lasso penalty in GL [23][22], is applied to the off-diagonal entries of the K precision matrices to encourage sparsity. The second term, which is the fused lasso penalty [28], is applied to the differences between elements of each pair of precision matrices to encourage similarity between conditions. A large λ_2_ leads to similar edge patterns across conditions. It has been shown that FGL outperforms GL when conditions are related [6].

### Condition-adaptive Fused Graphical Lasso

While borrowing strength across conditions is helpful for enlarging effective sample sizes, differences in co-expression patterns are present between different conditions. FGL encourages similarities among all edges across all conditions equally by imposing a constant penalty parameter for the fused lasso penalty. This has two drawbacks. First, it does not distinguish between shared edges and those unique to a condition, thus condition-specificity of edges is not preserved. Second, it encourages an equal amount of similarity across conditions without considering distance between conditions. For example, if one were studying tumor-specific co-expression by analyzing two subtypes of tumor tissues and a normal tissue jointly, FGL would impose an equal amount of similarities across all the pairs regardless whether the pair consists of two tumor subtypes or a tumor tissue and the normal tissue. This is problematic as the two tumor subtypes are likely to be more similar to each other than to the normal tissue.

To address these issues, we extend the fused graphical lasso method to incorporate condition-specificity in the integration of networks across conditions. Our strategy is to add a binary screening matrix **w**^(kk′)^ to the fused lasso penalty as follows,

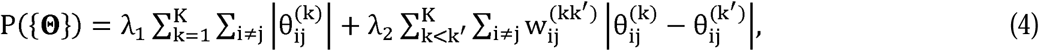

where 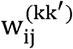 is the (i,j)-th element of **w**^(kk′)^ with

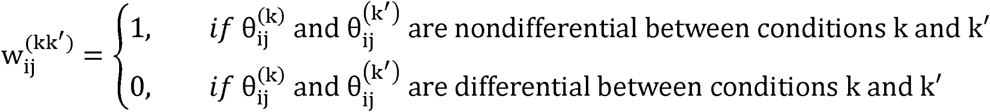

The matrix **w**^(kk′)^ controls whether similarity should or should not be encouraged between each pair of condition for each edge. It allows different edges to be penalized differently, and also allows the penalties for different pairs of conditions to vary according to the distance between the conditions. In doing so, one can borrow strength across conditions for estimating common edges, while allowing differential edges to be estimated in a condition-specific way. Therefore, we call our method condition-adaptive fused graphical lasso (CFGL).

### Determine the screening matrix **w**^(kk′)^

The screening matrix can be obtained using prior knowledge, learning directly from the data, or a combination of both strategies. To determine the screening matrix using prior knowledge, one may extract information on co-expression regulation from public databases, such as the KEGG pathway database [29] or COXPRESdb [30]. For example, if a pathway is known to be conserved across tissues [5][31], one may specify the corresponding elements in **w**^(kk′)^ as 1 to reflect the conservation of co-expression regulation.

Since prior information is often very limited or unavailable for many applications, we propose a data-driven strategy to estimate the screening matrix **w**^(kk′)^ from the data. As **w**^(kk′)^ reflects the status of differentiation between a pair of conditions, we determine **w**^(kk′)^ by identifying differential entries between the precision matrices of the two conditions, 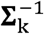 and 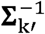, through a hypothesis test. If the test determines that the entry *ij* is differential, we set 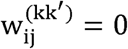, otherwise we set 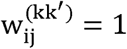. As 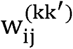 is binary, this approach is equivalent to using a l_0_ penalty to determine the support of the condition-specific edges. It is somewhat analogous to the Sure Independent Screening procedure for feature selection methods such as the lasso, Dantzig selector, and SCAD.[32], where an elementwise screening is first performed to reduce the dimension from ultra-high to moderate before variable selection.

Here we test for differentiation using the test proposed by Xia *et* al.[33]. This method tests for a difference between a pair of precision matrices and reports differential entries in the precision matrices with a proper false discovery rate (FDR) control. It directly estimates the difference between precision matrices, bypassing the estimation of the individual precision matrices. Other tests for differential entries are available[34][35], but we selected Xia’s test as it has been shown to provide more accurate estimates than the tests that require separate estimation of precision matrices due to leveraging information on the sparsity of the difference between precision matrices [36]. To avoid falsely imposing similarity for edges that are moderately differential, we use a relaxed FDR threshold in the test to encourage similarity only to the edges that are obviously non-differential across conditions.

### Parameter estimation and selection of penalty parameters

Similar to FGL and other penalty-based methods, this model can be estimated using the ADMM algorithm. We used BIC to guide the selection of penalty parameters. In the real data application, when the sample size is reasonably large to afford subsampling, we performed an additional stability selection [37] step. Instead of constructing networks using all the samples, the stability selection procedure constructs networks for a large set of subsamples generated from the original data and keeps only the edges that frequently occur across subsamples to obtain robust edges. Details on the stability selection procedure can be found in Methods.

## Simulation studies

We used simulation studies to evaluate the performance of our method and compare it to FGL and GL. We first considered the two-condition scenario, evaluating the performance of these methods at different levels of differentiation between the conditions. Then, we increased the complexity by introducing a third condition and allowing the level of differentiation to vary across all three conditions.

In the first set of simulations, we generated the gene expression profiles from a co-expression network of 400 genes for 2 conditions. The network consists of 8 co-expression modules, each of 50 genes. To generate different levels of differentiation between conditions, we simulated four scenarios (S1-S4) with a progressively increasing number of differential edges between conditions, where the networks in the two conditions are identical in S1 and are different at various levels in S2-S4. While S1 is extremely rare in practice, it exactly follows the model assumptions of FGL and thus illustrates the methods performance under conditions ideal to FGL. Two samples size (50, 100) are considered for each scenario.

To simulate a network, we first simulated its constituent modules. To create different levels of differentiation, three types of modules were simulated: (1) identical network structure and identical edge weights between conditions (II), (2) identical network structure but different edge weights between conditions (ID), and (3) different network structure and different edge weights between conditions (DD). We then combined these modules in various configurations to achieve the desired level of differentiation for the networks in different scenarios. In all scenarios, the 8 modules are evenly split into two groups, each of which consists of 4 modules of the same type. The configurations of modules in these scenarios are summarized in Table 1. Detailed information on the data generating process are in Methods.

**Table 1.**
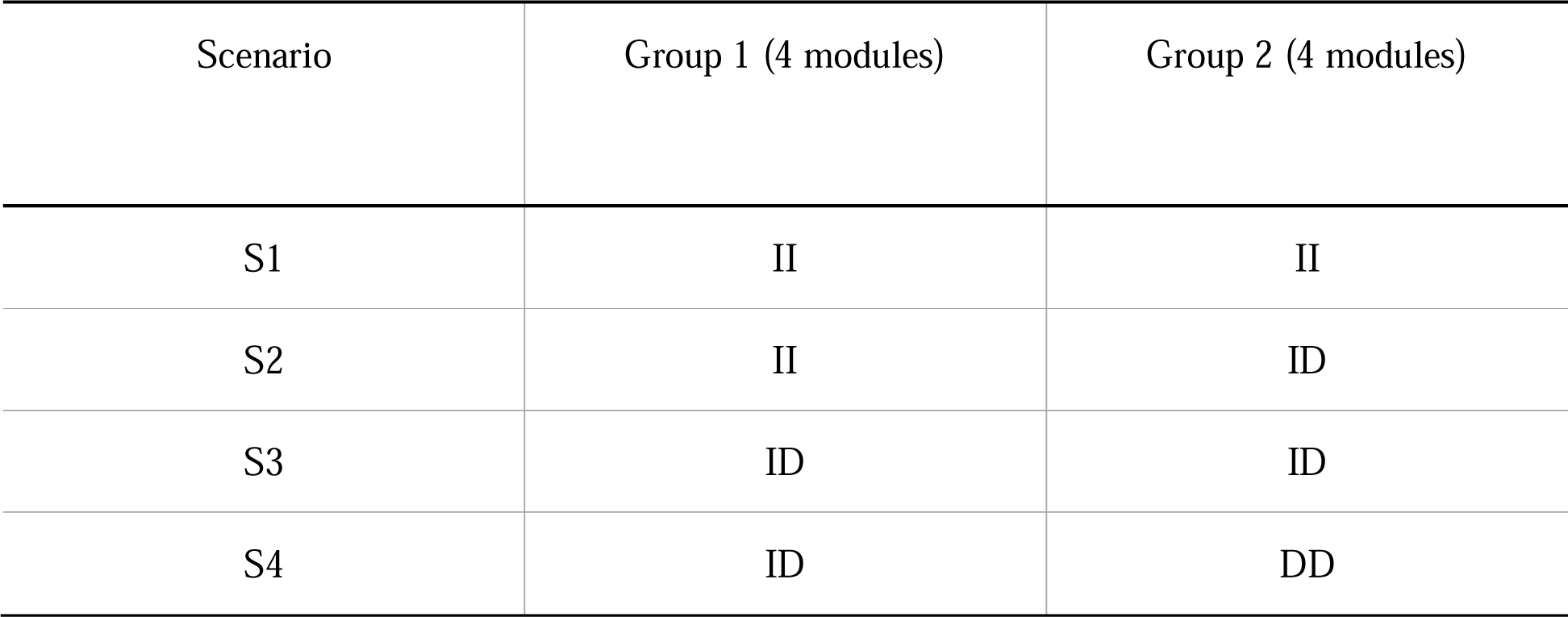
Configurations of simulation scenarios. The level of differentiation between two conditions in the constituent modules is shown in Columns 2-3. II: identical network structure and identical edge weight; ID: identical network structure and different edge weights; DD: different network structures and different edge weights.

| Scenario | Group 1 (4 modules) | Group 2 (4 modules) |
| --- | --- | --- |
| S1 | II | II |
| S2 | II | ID |
| S3 | ID | ID |
| S4 | ID | DD |

For each simulation, we constructed the co-expression network using our method, FGL, and GL. To evaluate how the accuracy of the estimated screening matrix affects the performance of our method, we also included a version of CFGL with the true screening matrix, which is labeled as CFGL-oracle (CFGLO) (see the Methods section). We compared the performance of these methods in terms of edge detection and edge weight estimation. To evaluate the accuracy of edge detection, we computed the true positive and false positive rates across a grid of *λ*_1_ and *λ*_2_. The accuracy of edge weight estimation was assessed by computing the sum of squared error (SSE) between the estimated edge weight and the true precision matrix. We plotted the ROC curve for edge detection and the SSE for edge weight estimation at a varying level of *λ*_1_ with *λ*_2_ fixed at the value that achieves the minimal BIC value (*λ*_2_ = 0.15 for *n* = 50 and *λ*_2_ = 0.10 for *n* = 100).

Fig 1 shows the results at *n* = 50. In all scenarios, our approach (CFGL) had a higher AUC and a lower SSE than GL. The gain is more apparent when the two conditions are relatively similar (S1-S3). This is because data integration improves the accuracy of edge detection, especially when networks are similar between conditions. The advantage is especially obvious when n=50 (Supplementary Table 1 and Supplementary Fig 1 for n=100), as this small sample size is likely not enough to support accurate estimation with GL based on the samples from a single condition. FGL performs well in these scenarios too; however, when the two conditions are fairly different (S4), FGL performs worse than GL. Compared with FGL, our method has a higher AUC and an apparently lower SSE in all scenarios with between-condition differences (S2-S4). Even in the scenario without between-condition differences (S1), i.e. the ideal setting for FGL, our method is still competitive: it has an almost identical ROC curve as FGL and a slightly higher SSE than FGL. In practice, it is much more common to encounter S2-S4 than S1, as the networks of two different conditions are likely to be different. CFGLO has the best AUC and SSE among all the methods, suggesting that the performance of CFGL can be further improved by improving the estimation of screening matrix, for example, by incorporating external information. We also reported simulation results under several other *λ*_2_. The results are similar and can be found in Supplementary Table 1.

**Fig 1.**
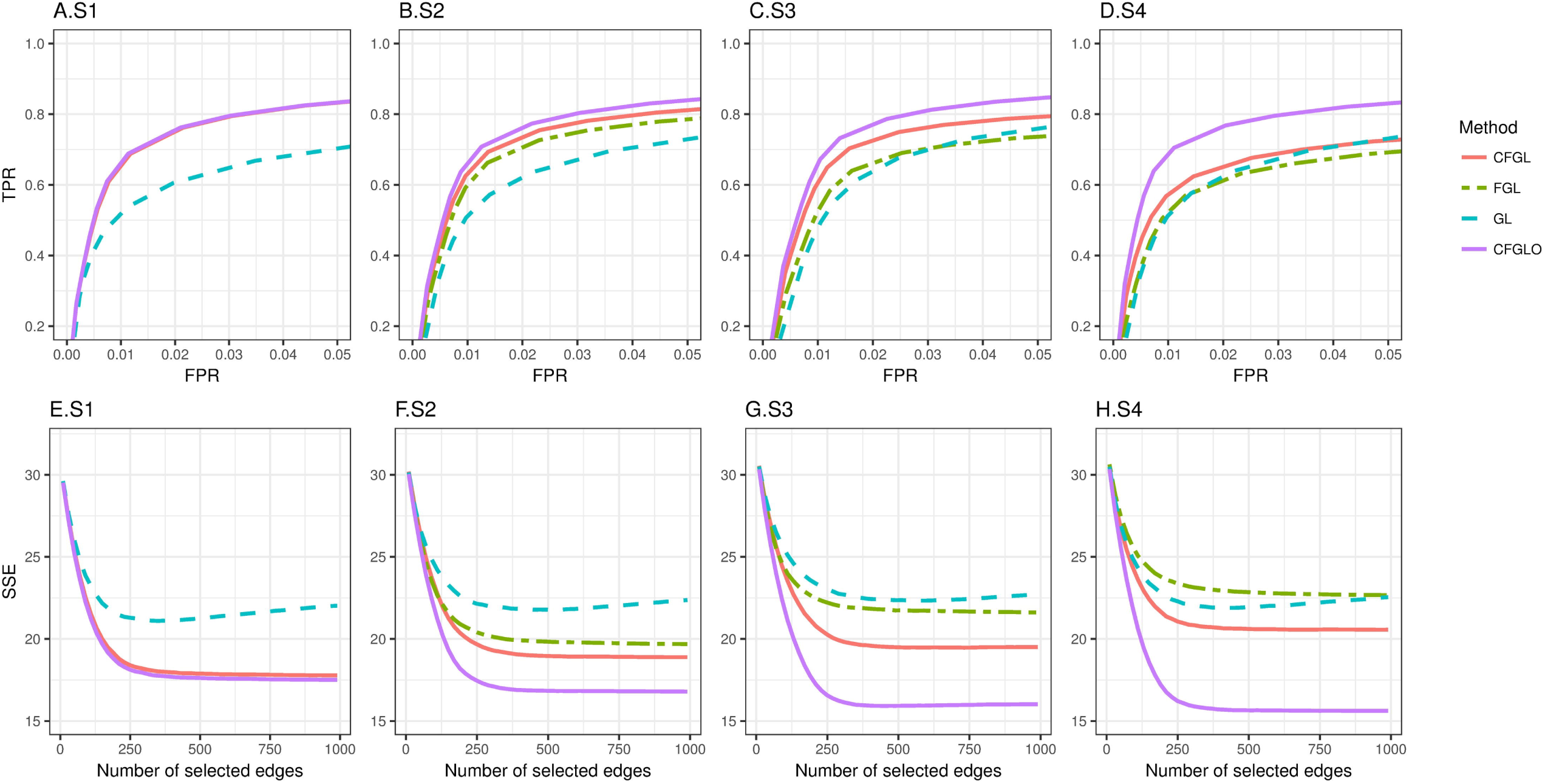
Performance comparison for simulations with two conditions. Top row: ROC curves for edge detection. Bottom row: SSE for edge weights estimation. Red line: CFGL, Green line: FGL, Blue line: GL, Purple line: CFGL-oracle.

Next, we allowed the level of differentiation between conditions to vary across conditions. Such a situation commonly arises when one performs co-expression network analysis for multiple conditions. Here, we simulated the gene expression profiles under three conditions for 450 genes comprised of 9 modules of 50 genes each. Similar to the 2-condition simulation, we included two groups of 4 modules of the same type. To better imitate real networks, we also included an additional type II module to mimic housekeeping co-expression across 3 conditions. In total, we considered 4 scenarios. Table 2 summarizes the configurations of these simulations. S1 and S2 represent the cases where pairwise similarities between conditions are constant across conditions, with a higher similarity in S1 than in S2. S3 and S4 represent the case where pairwise similarities vary across conditions, with a higher similarity in S3 than S4. In this simulation, we compared CFGL, FGL and GL. CFGLO was not included as its performance is similar to the previous case.

**Table 2.**
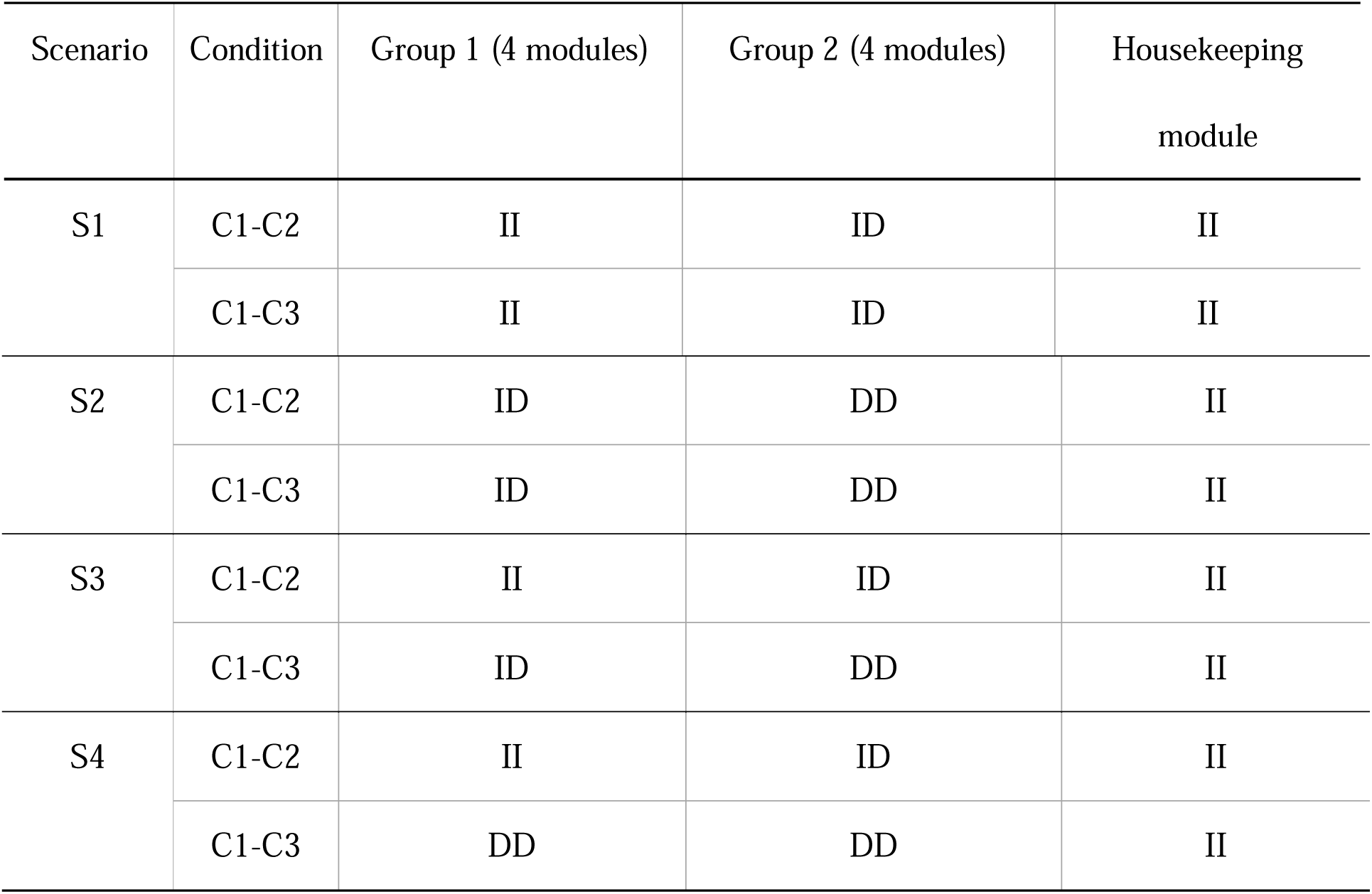
Configurations of scenarios in the 3-condition simulation. The pairwise similarity between condition 1 and other conditions is reported.

In all scenarios, our approach has a higher AUC and a lower SSE than both GL and FGL (Fig 2, Supplementary Table 2). The gain over GL is most apparent when the differentiation between conditions is low (S1). This is again because data integration is most beneficial when networks are similar across conditions. FGL performs well in this case too. However, when the distance between conditions is different across conditions (S3), the advantage of FGL over GL diminishes; and when the differentiation between conditions is relatively high (S2 and S4), FGL performs worse than GL. This is expected, as imposing similarity across conditions as in FGL is improper for these scenarios. However, our method performs well in all scenarios.

**Fig 2.**
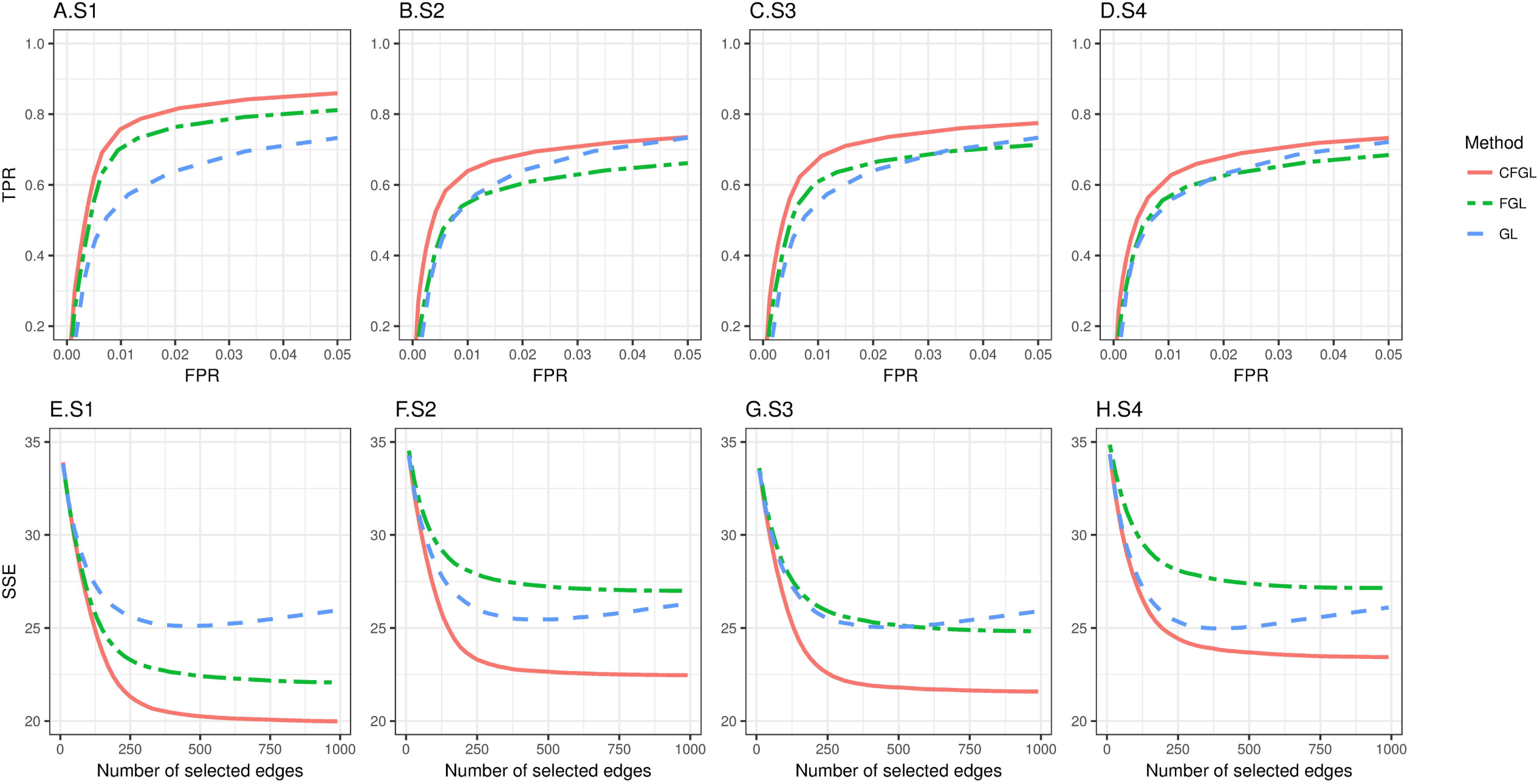
Performance comparison for simulation with 3 conditions. Top row: ROC curve for edge detection (A-D) Bottom row: SSE for edge weights estimation (E-H). Red line: CFGL, Green line: FGL, Blue line: GL.

Taken together, we attribute the gain of our methods to its adaptive way of enforcing similarities. When networks are highly similar across conditions, enforcing similarities across all edges, as in FGL, is optimal. Our method adapts to this situation and produces similar results to FGL. In contrast, when networks are different across conditions, similarity should be encouraged only among the shared edges in data integration. Our method is again adaptive to the differential patterns across conditions, thus shows more gain when the difference between conditions is present.

### Application to rat expression data

We applied our method to a microarray dataset collected from a recombinant inbred (RI) rat panel and compared with FGL and GL. The gene expression profiles in the brain and heart tissues were measured for 19 rat strains using Affymetrix Rat Exon Array 1.0 ST. Details on data processing and normalization are provided in Methods. Because of the small sample size, we restricted the network construction to the 500 most differentially expressed (DE) genes between brain and heart (see Methods), and used BIC to guide the selection of penalty parameters.

#### Tissue specificity for edges

Since brain and heart have different embryonic origins and functions, a considerable number of tissue-specific co-expression relationships are expected. We first compared the co-expression networks constructed by each method in terms of tissue specificity (Supplementary Fig 2-3). Fig 3A shows the number of edges identified by each method at the optimal BIC (*λ*_1_ =0.0010 and *λ*_2_ =0.0008 for both CFGL and FGL, and *λ*_1_ =0.0009 for GL), categorized by their tissue specificity. Our method and FGL identify substantially more tissue-common edges than GL. For example, at *λ*_2_ = 0.0008 (Fig 3A), 26.0% (356 out of 1374) and 35.6% (354 out of 994) of edges detected by our method and FGL, respectively, are common between tissues; whereas only 0.2% (3 out of 1491) of edges detected by GL are common between tissues (Supplementary Table 3). This is expected, as the fused penalty in CFGL and FGL enforces similarities across tissues. Compared with FGL, our method detects substantially more tissue-specific edges. To check if this pattern is related to the choices of *λ*_2_, we also performed the same analysis at *λ*_2_ =0.0010 and 0.0012 (Fig 3 B-C). This observation persists at all *λ*_2_ levels. This difference is especially obvious when a relatively high *λ*_2_ is applied: the proportion of tissue-specific edges detected by FGL rapidly reduces, whereas our method still maintains a considerable proportion of tissue-specific edges. For example, when *λ*_2_ =0.0012 (Fig 3C), our method detects 23.6% (191) brain-specific edges and 20.6% (161) heart-specific edges, whereas FGL detects only 1.1% (7) brain-specific edges and no heart-specific edges.

**Table 3.**
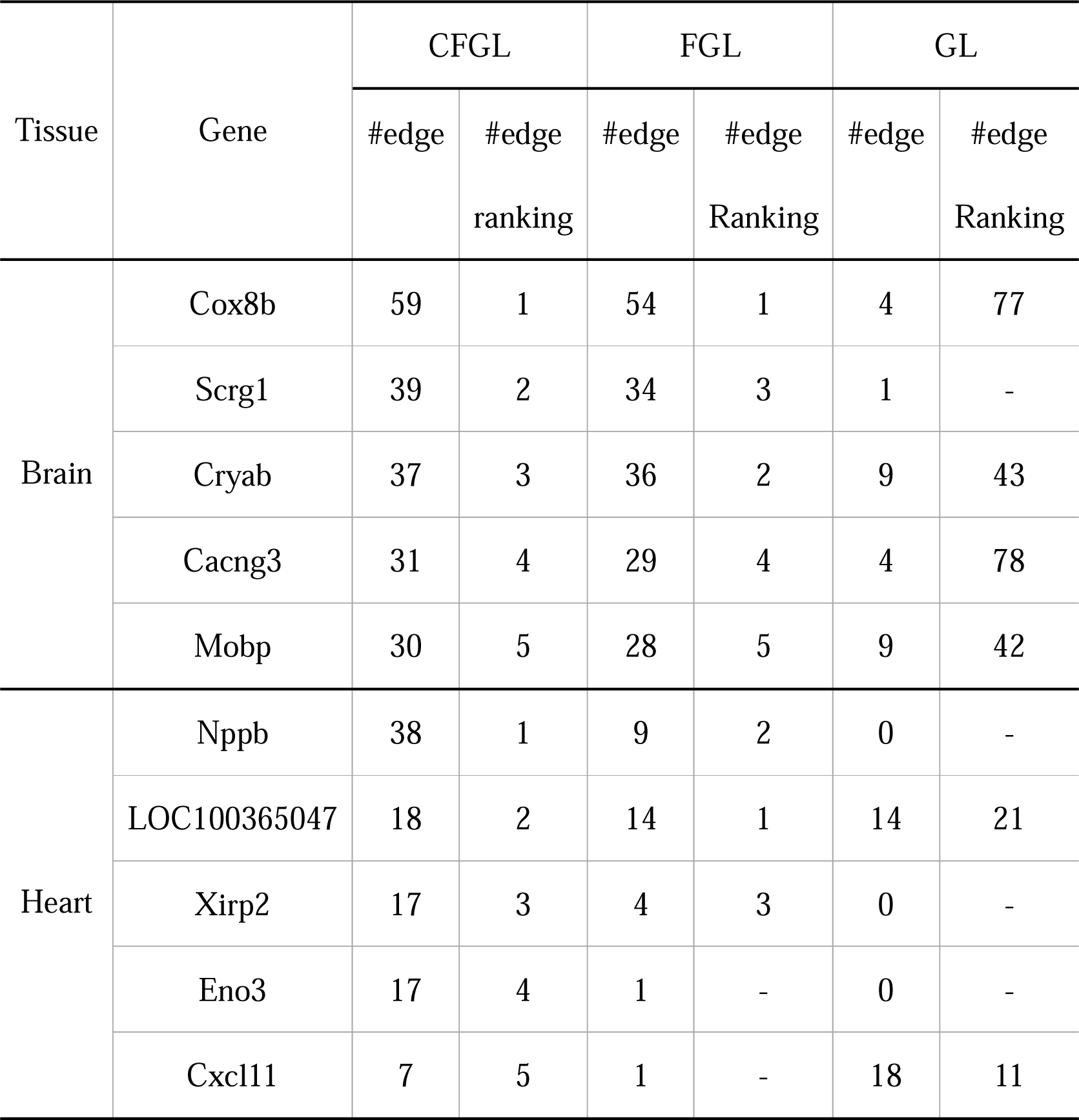
Top-5 tissue-specific hub genes identified by our method. The numbers of tissue-specific edges linked to the hub and the corresponding rankings are reported, in comparison with the results from FGL and GL.

**Fig 3.**
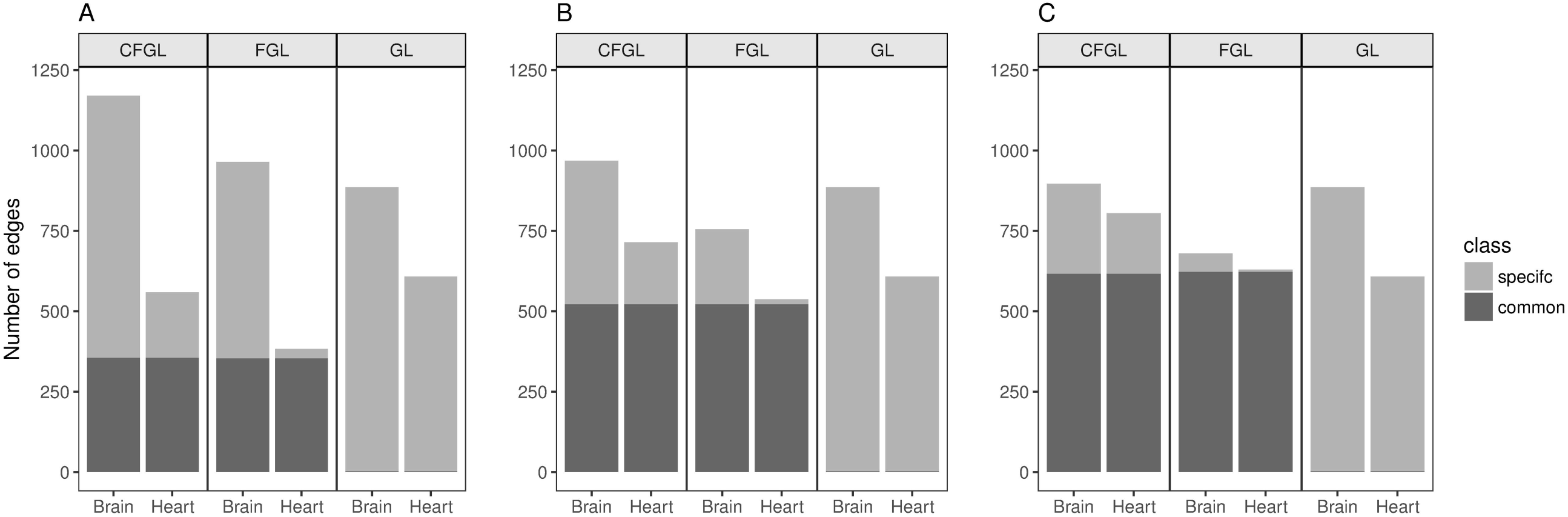
The numbers of tissue-specific and tissue-common edges detected by our method, GL, and FGL.

#### Tissue-specific hub genes identified by CFGL demonstrate highly tissue-specific biological functions

Because the estimated heart- and brain-specific networks are both considerably dense, it is difficult to identify disjoint co-expression modules. Instead, we first identified tissue-specific hub genes, and then formed a tissue-specific module for each hub gene using the hub gene and the genes directly connected with the hub by tissue-specific edges. To identify these hub genes, we counted the number of edges that are specific to each tissue for each gene and reported the five genes with the highest number of tissue-specific edges identified by our method (at *λ*_1 =_ 0.001, *λ*_2 =_ 0.0008) in Table 3. Because the genes used to construct networks in this analysis are differentially expressed genes, any random set of them presumably would show some tissue specificity. However, a random set of DE genes is much less likely to form a functionally coherent module than a set of co-expressed DE genes. Thus the functional coherence of a module helps establish confidence in the identified co-expression pattern. To examine the functional coherence for these modules, we performed a Gene Ontology (GO) enrichment analysis using the non-hub genes in each module and checked its agreement with the functionality of the hub gene. As a comparison, we also performed a GO enrichment analysis using a set of the most differentially expressed genes with the same size as that of the identified module.

Among the brain-specific hub genes, Mobp has been found to be specifically expressed in oligodendrocytes. It encodes the protein related to sheath compaction in rat brain and spinal cord [38]. Its peripheral genes also show a significant GO enrichment in myelin sheath (FDR=1.558E-3), consistent with the function of Mobp. Interestingly, this GO term was not enriched using the set of 30 most differentially expressed genes in brain, providing an example of how differential co-expression analysis can uncover biologically important findings not revealed by differential expression analysis alone. Cacng3 is a protein-coding gene that encodes type I trans-membrane AMPA (α-amino-3-hydroxy-5-methyl-4-isoxazolepropionic acid) receptor regulatory protein (TARP). Its product, TARP gamma-3, is abundant in the cerebral cortex and amygdala, and has been found to be associated with childhood absence epilepsy in humans [39][40]. The genes specifically co-expressed with Cacng3 in brain are also enriched in the GO terms related to neuron differentiation (FDR=1.332E-2), anterograde trans-synaptic signaling (FDR=1.332E-2) and synapse (FDR=4.161E-6). Among heart-specific hub genes, Nppb (also known as BNP) is a member of the natriuretic peptide family and encodes a secreted protein that functions as a cardiac hormone. It has been reported that Nppb is associated with an intra-cardiac counterregulatory mechanism that prevents the development of cardiac fibrosis in vivo. It has been suggested that this gene serves as a local regulator during the process of ventricular remodeling [41]. The genes specifically connected with Nppb in heart are also enriched in the GO terms related to cardiac muscle tissue development (FDR=3.886E-3). Another heart-specific hub gene, Xirp2 (also known as CMYA3), has been reported to be related to the formation of intercalated disc (ICD), which is a juncture that links cardiac muscle cells and plays vital roles in signaling among cardiomyocytes[42]. The genes connected to Xirp2 are also enriched in GO terms related to muscle fiber development (FDR=1.517E-2).

FGL identifies the same set of brain-specific hubs and a similar brain-specific subnetwork. However, our method detects more heart-specific hubs than FGL, and each hub harbors more heart-specific edges (Table 2). GL reports drastically different results (Supplementary Table 4) from FGL and our method in both tissues. None of the hubs in Table 2 are ranked among the top 5 by GL.

### Application to TCGA data set

We applied our method to the breast cancer data from the TCGA project. Breast cancer is the most common cancer among women [43]. According to the presence and absence of the estrogen receptor (ER) in cancer cells, breast cancer can be classified into two subtypes, ER+ and ER-. Approximately two-thirds of breast cancer are ER+ at the time of diagnosis, and the rest are ER-. The ER status provides important clinical implications for both mechanisms of carcinogenesis and therapeutic treatment [44].

The TCGA project [43] has collected gene expression RNA-seq data for 1100 breast cancer patients. Among them, 112 individuals have both tumor tissue and matched peripheral normal tissue. Our goal is to identify co-expression modules that are specific to ER+ or ER-subtype and those that are shared between the two tumor subtypes but are not present in normal tissue. To ensure the independence of the samples in our analysis, we used the normal tissue samples from these 112 individuals and tumor samples that are annotated as ER+ (187 samples) or ER- (98 samples) from different individuals. Due to the limited sample size, we restrict our analysis to a subset of 1000 genes that either show a significant association with survival time in a Cox model or have been reported to be related to breast cancer (Details in Methods). To obtain robust co-expression networks, we applied the stability selection procedure in [37] to CFGL, FGL, and GL. Details of the data processing steps and the procedure of the stability selection can be found in Methods.

#### Disease type specificity of the co-expression edges

To investigate disease type specificity of the edges, we partitioned the identified edges into seven mutually exclusive categories: normal tissue only, ER+ subtype only, ER-subtype only, normal and ER+ shared, normal and ER- shared, ER+ and ER- shared, and all tissue common. The number of edges identified by each method in each category is summarized in Supplementary Table 5. Fig 4 compared the disease type specificity of the edges identified by the three methods. Similar to the rat dataset, the majority of the edges identified by GL are unique to one tissue with a very small percentage (68/5978=1.1%) of edges shared by all tissues. In contrast, a high percentage of edges identified by FGL (1330/4448=30.0%) and our method (684/2624=26.1%) are common across all tissues.

**Fig 4.**
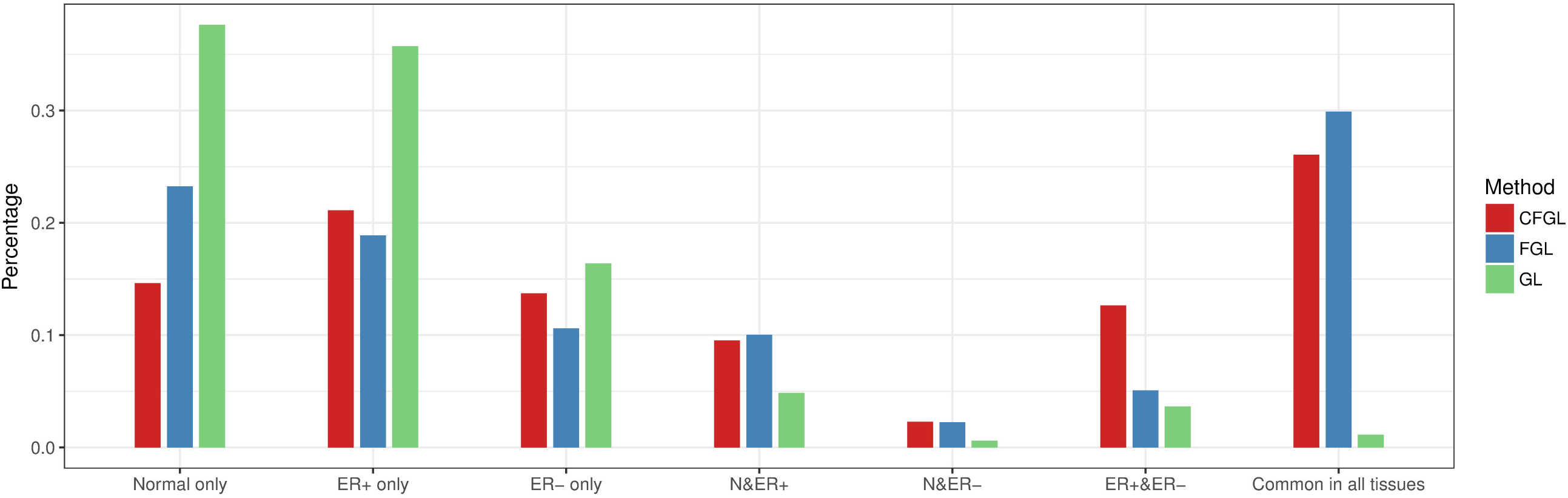
Disease type specificity of the estimated co-expression edges for the TCGA data.

We also evaluated the pairwise similarity of the co-expression networks between each pair of tissues (Fig 4). As ER+ and ER-tumors both are subtypes of breast cancer, they are expected to be more similar to each other than to normal tissue. However, we observed that FGL and GL show a higher proportion of shared edges between normal tissue and ER+ tumor (FGL: 446/4448=10.0%; GL: 290/5978=4.9%) than between the two tumor subtypes (FGL: 226/4448=5.1%; GL: 218/5978=3.6%). In contrast, our method shows a much higher proportion of shared edges between the two tumor subtypes (332/2624=12.6%) than between the normal tissue and either of the tumor subtypes (normal and ER+: 250/2624=9.5%; normal and ER-: 60/2624=2.3%), reflecting the expected biological similarity. The proportion of common edges between the two tumor subtypes is also substantially higher in the network constructed by our method than in those constructed by the other two methods (CFGL: 332 (12.6%), FGL: 226 (5.1%), GL: 218 (3.6%)).

#### Genes most significantly associated with survival time usually are not hubs

To investigate the biological relevance of the hub genes in the co-expression network constructed by our method, we examined the relationship between the hubness of each gene, i.e. the number of edges connected to the gene, and its association with survival time. Strikingly, we found a clear negative correlation between the hubness of a gene and the significance of its association with survival time (Fig 5): the genes that have the most significant p-values in the Cox model usually are not hub genes, whereas hub genes tend to have less significant p-values. One possible explanation is that many genes that are significantly associated with survival time govern very particular functions, thus they are not correlated with the expression of many other genes. This agrees with the observation in the previous analysis [5] of the co-expression network for 35 human tissues in the GTEX project, in which genes with tissue-specific functions were observed to have fewer co-expression edges than average.

**Fig 5.**
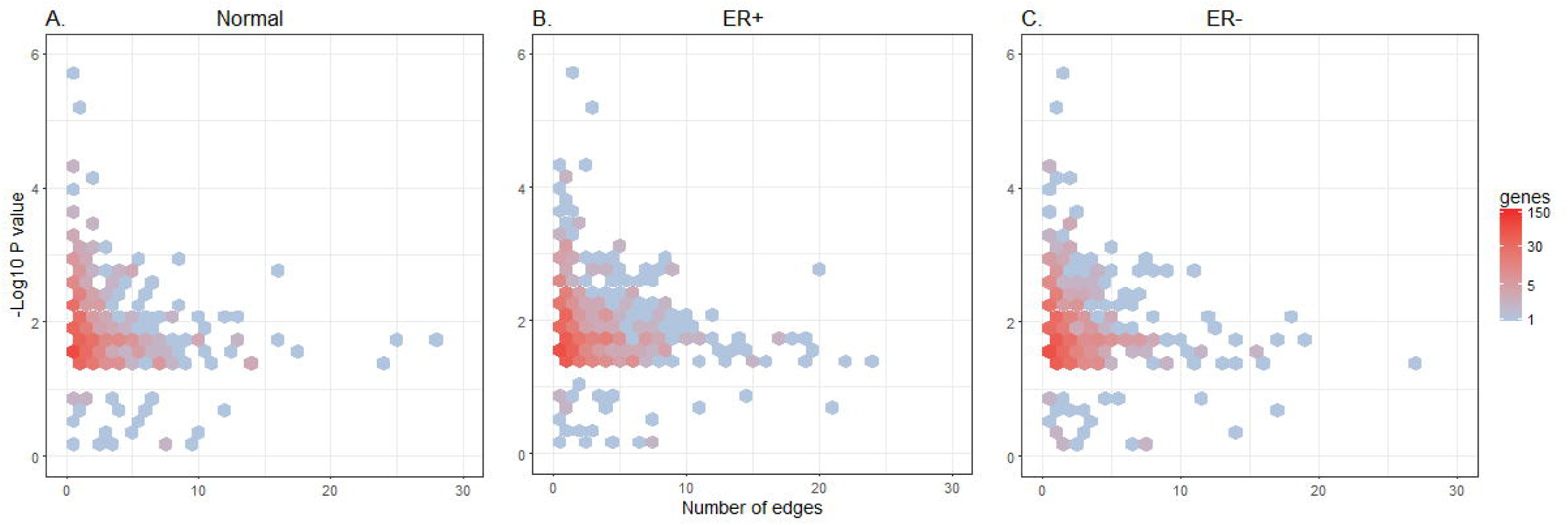
Hubness of a gene and its association with survival time in three tissues. Y-axis: -log10(p-value) of a gene in the Cox model. X-axis: the number of edges connected to a gene. The gene set consists of 961 genes that are significant associated with survival time (p-values <0.05) and 39 genes that are not significant but are known to be related to breast cancer (see Methods).

#### Hubs specific to a disease type tend to not share edges across all tissues

To understand the role that a gene plays in condition-specific and condition-common co-expression, we classified edges according to their disease type specificity and compared the hubness of a gene in the network shared by all three tissues (3T) and the network specific to one tissue (1T). We observed a clear negative correlation between these two types of hubness (Fig 6), especially in the two tumor tissues. For all genes with at least 5 edges, the Spearman correlation between the numbers of 1T edges and the numbers of 3T edges is -0.23, -0.69, and -0.43 in normal, ER+ tumor, and ER-tumor tissue, respectively. It can be clearly seen that several ER+ or ER-specific hubs have very few edges shared across all tissues. This indicates that co-expression hubs specifically triggered in tumor tissues are usually not the co-expression hubs in normal tissue. These hubs may provide important insights in carcinogenesis and cancer treatment. The tumor-specific hubs and their possible biological functions are discussed in the next sections.

**Fig 6.**
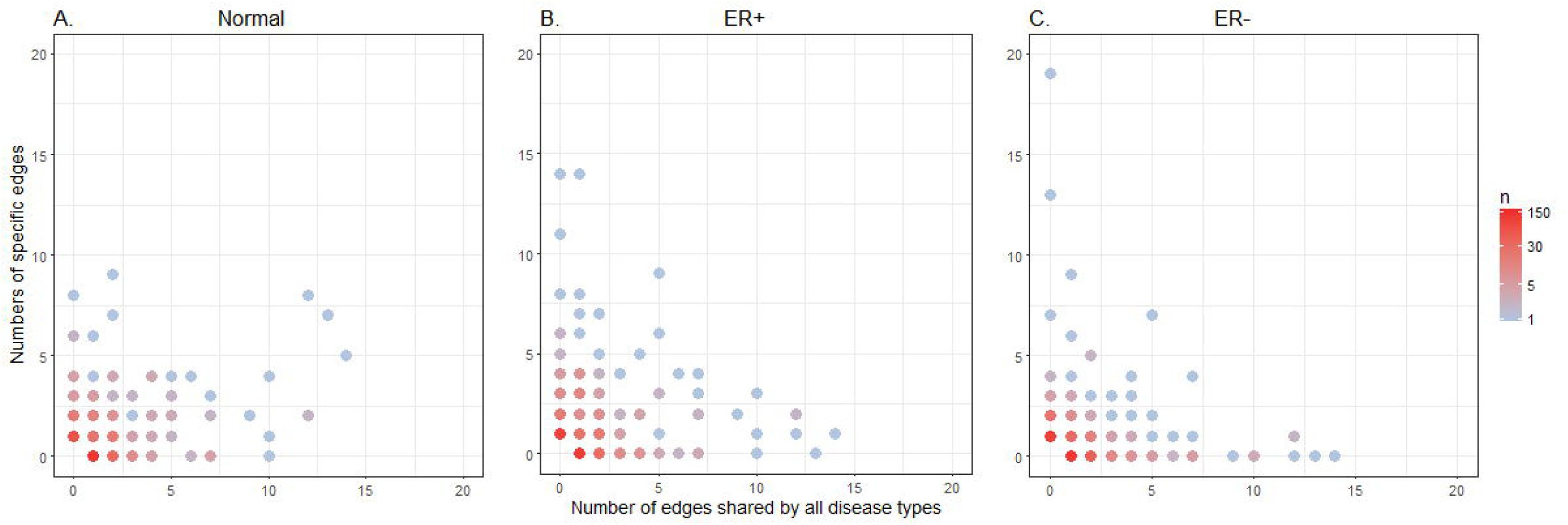
Distribution of disease type-specific edges and -common edges for genes in the co-expression network constructed using our method.

#### Biological functions of tumor-related modules

Next, we characterized the subnetwork shared between ER+/ER-, ER+ specific subnetwork and the ER-specific subnetwork in order to further study the biological function of the co-expression network of tumor tissues. We obtained each subnetwork by extracting the corresponding disease type-specific edges from the networks constructed using our method. For each subnetwork, we identified major disjoint modules and then annotated their biological functions using GO enrichment analysis. To determine the hub gene in each module, we counted the number of edges for each node in the module (Table 5). For ER+ specific and ER-specific modules, we characterized all genes that have more than 10 edges. Because the size of tumor-shared modules is relatively small, we only characterized the genes with the most edges in the module.

**Table 5.**
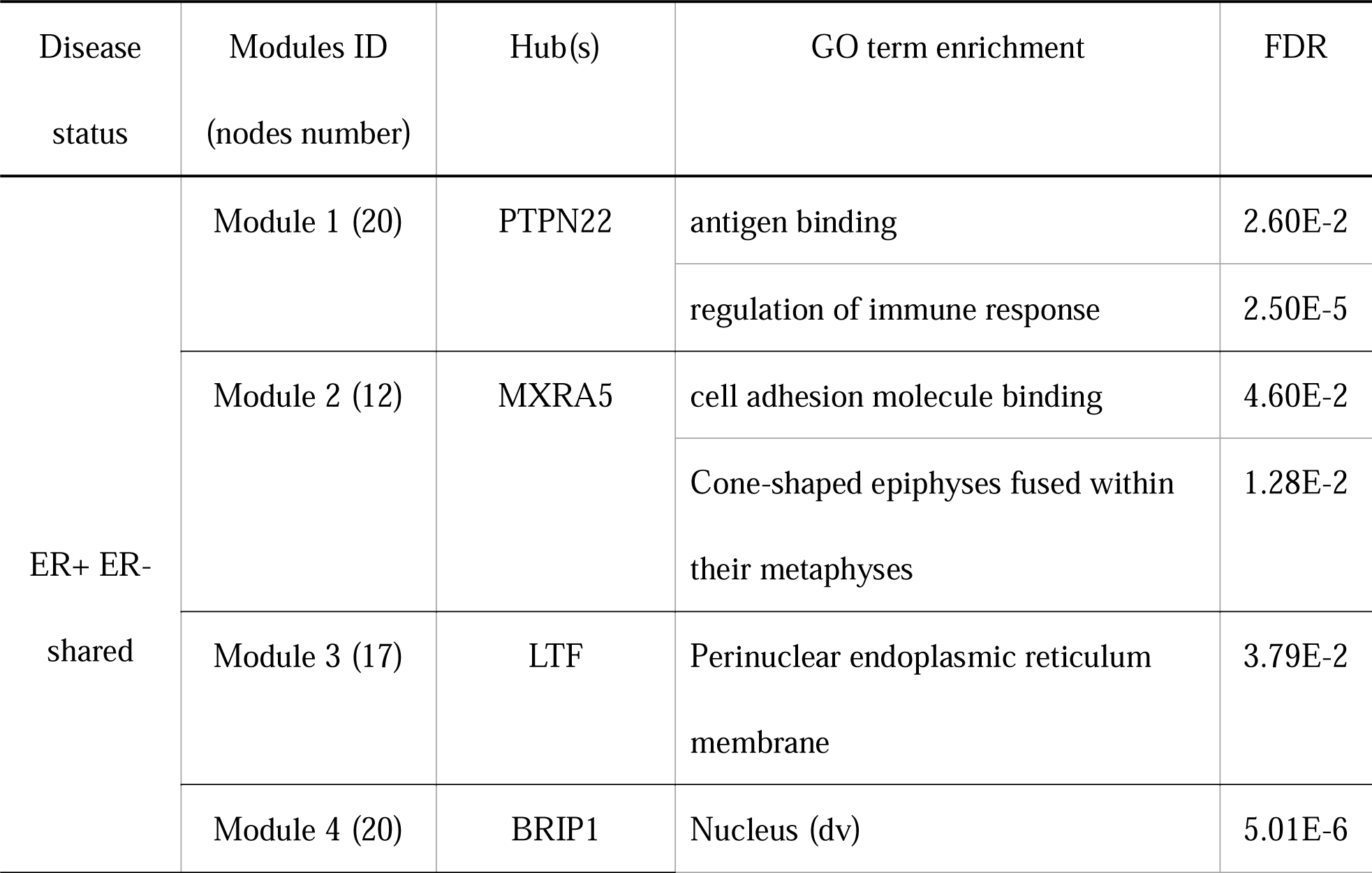
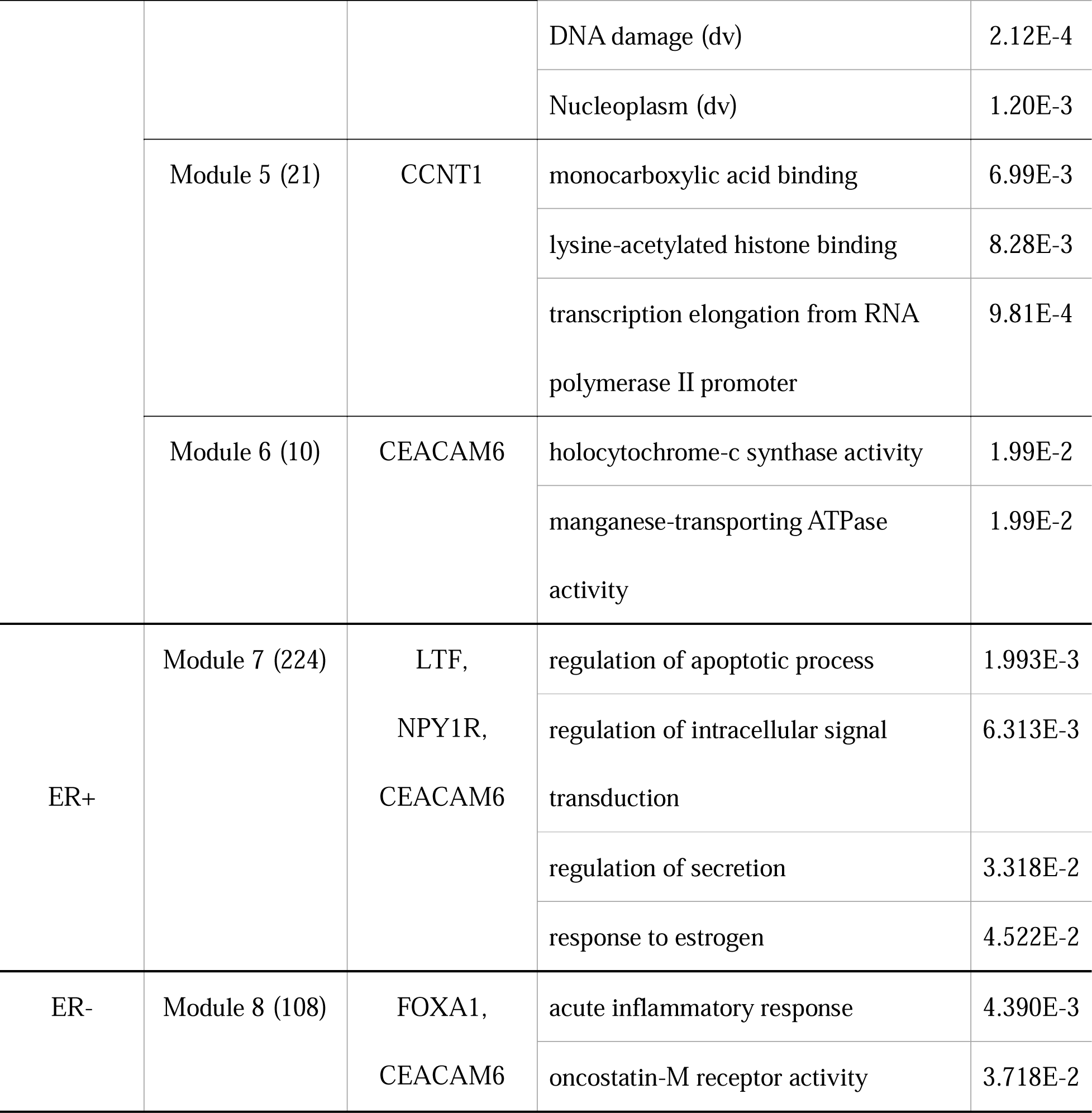
Modules and their significantly enriched GO terms.

#### Tumor-shared subnetwork

The tumor-shared subnetwork consists of 203 genes and 332 co-expression edges (Supplementary Fig 4). There are 6 disjoint co-expression modules with more than 10 genes (Fig S1). All 6 modules show significant enrichment (FDR<0.05) in the GO term analysis. They are enriched in GO terms of Immunity, antigen processing and presentation, Nucleus, and DNA damage (Table 4).

Among the hubs of the 6 modules, 5 of them (PTPN22, BRIP1, CEACAM6, LTF, and CCNT1) have been previously found to be associated with breast cancer [45][46][47][48][49][50][51][52], and the other one, MXRA5, has been reported to be related to non-small cell lung cancer[53][54]. PTPN22 encodes a protein tyrosine phosphatase that is involved in the signaling pathways associated with immune response (Fig 7). Previous studies have shown that overexpression of PTPN22 significantly inhibits the growth of human breast cancer cells. Its product also blocks cancer cell xenografts and their metastases[45]. BRIP1 (also known as BACH1, FANCJ), together with BRCA1, is involved in the repair process of DNA double-strand breaks. It has been reported that BRIP1 acts as a master regulator of breast cancer [55]. Previous studies have reported that overexpression of BRIP1 promotes the migration and invasion of cancer cells, while knockdown of BRIP1 suppresses this process[56]. ZKSCAN1 (also known as KOX18, ZNF139), a node with three connections in the same module with BRIP1, also has been reported to play regulatory roles in migration and invasion of human gastric cancer cells [57][58]. CEACAM6 encodes a protein in the carcinoembryonic antigen family, which has been shown to be associated with cell adhesion. Previous studies have shown that CEACAM6 is detected in approximately 70% of solid tumors, including breast cancer [43][59]. It has been suggested that CEACAM6 is associated with tumor progression stage [60], inhibition of cell differentiation and anoikis, and promotion of cell adhesion, invasion, and metastasis [47]. LTF was previously found to inhibit the growth of solid tumors and the development of experimental metastases[48][49][50]. Overexpression of CCNT1 was found as an implication of tumor growth[52].

**Fig 7.**
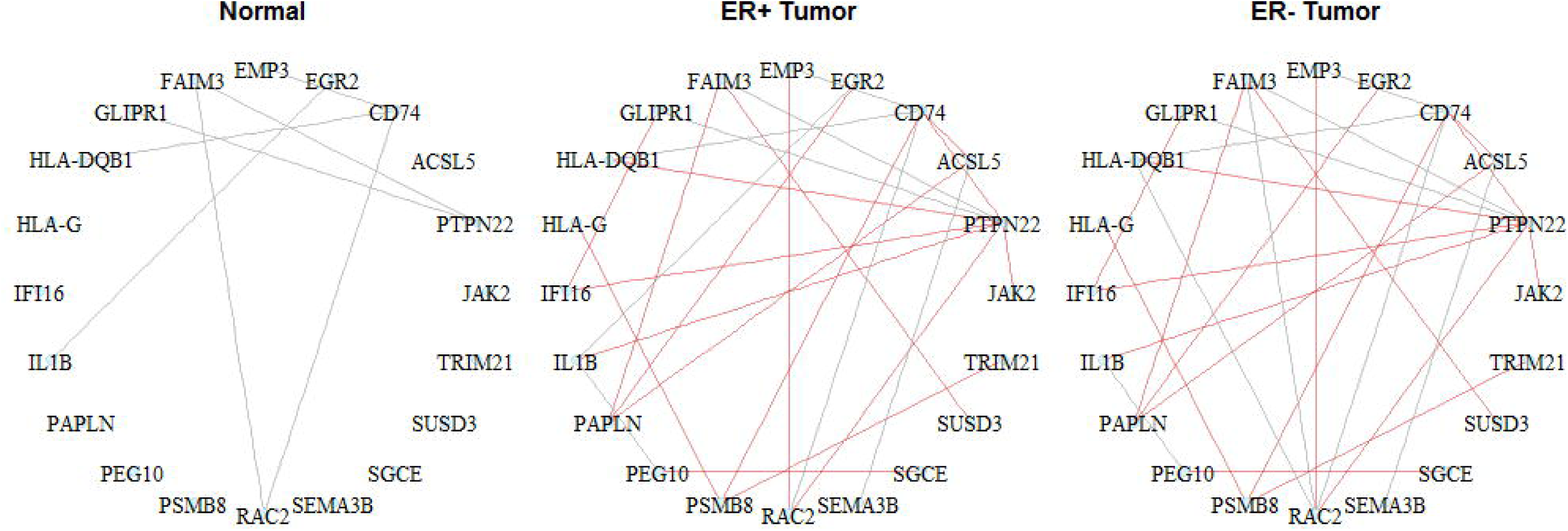
An example of ER+/ER-shared module in TCGA breast cancer data. The genes in the module are enriched in GO term of Antigen binding. The hub gene of the module, PTPN22, was found associated with an immune function in breast cancer. Red: common edges shared by two tumor tissues. Grey: all other edges.

#### ER+ specific subnetwork

The ER+ specific subnetwork consists of 554 edges and 296 nodes (Supplementary Fig 5). One major co-expression module with 224 nodes is detected (Fig S2). Table 4 shows the enriched GO terms and the hub genes for this module. This module is enriched in the GO term related to the response to estrogen. One of its hub genes, NYP1R, has been reported to be involved in the activation of estrogen signaling pathway in breast carcinoma. It is up-regulated in ER+ tumor compared to ER-tumor [61][62]. This agrees well with the classification of ER+ subtype, which is characterized by the presence of estrogen receptors. The other two hub genes, LTF and CEACAM6, are also hubs genes in the tumor-shared modules.

#### ER-specific subnetwork

The ER-specific network consists of 360 edges and 223 nodes (Supplementary Fig 6). One major co-expression module with 108 nodes is detected (Fig S3). The GO term analysis shows that ER-specific module is enriched in the activity of the oncostatin-M receptor. This receptor is involved in the signaling event of oncostatin-M, a growth regulator that inhibits the proliferation of a number of tumor cell lines. Interestingly, we observed that, though CEACAM6 is a hub in both ER+ and ER-specific modules, it connects with different sets of genes in the two subtypes (Table 4). This indicates that CEACAM6, which is associated with tumor progression stage [60], may regulate cancer progression through different mechanisms in these two tumor subtypes.

### Conclusions

In this paper, we present a method, called condition-adaptive fused graphical lasso (CFGL), to construct gene co-expression networks for multiple conditions simultaneously. By incorporating a data-driven penalty that reflects the condition-specific co-expression pattern in the FGL framework, this method takes condition specificity into account while borrowing information across conditions in the network construction. It outperforms GL and FGL methods in both edge detection and estimation of edge weights across a range of scenarios in simulation studies.

Our analysis on a rat multi-tissue dataset and the TCGA breast cancer data reveals interesting biological insights. In both datasets, the modules in the condition-specific subnetwork identified by our method consistently show biologically relevant functions, demonstrating the suitability of our method for studying tissue-specific or disease-specific co-expression networks. The analysis on TCGA breast cancer data also reveals several interesting findings related to the mechanism of ER+ and ER-tumor subtypes. We found that the genes most significantly associated with survival time are less likely to be hubs. This suggests that most genes associated with cancer progression govern specific functions, rather than regulating a large number of biological processes. Similarly, we also observed that the hub genes in the tumor-specific subnetworks tend to not harbor edges shared with normal tissue. Several previously known cancer-related genes, including PTPN22, BRIP1, and CEACAM6, were found as hubs in the tumor-related subnetworks. Together, these results confirm the biological relevance of the results from our method. Other hubs and their specific co-expression edges are also worth further investigation.

Though our method was motivated by co-expression networks, it is suitable for any data application with multiple conditions but shared network structures, such as learning condition-specific binary networks with sparse Ising models [63][64]. The R package for our method CFGL is available on GitHub, https://github.com/Yafei611/CFGL.

### Method

#### Optimization of CFGL network

CFGL estimates the precision matrices {Θ} by solving

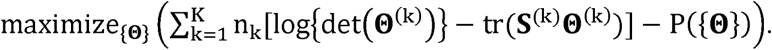

The penalty term p({Θ}) is

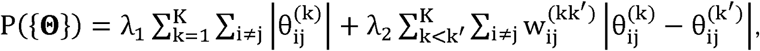

where n_k_ is the sample size of kth condition and **S**^(k)^ = (**y**^(k)^)^T^ **Y**^(k)^/n_k_ is the empirical covariance matrixes for the kth expression data set.

We implemented the ADMM algorithm [65] to solve the above problem. The detailed optimization procedure can be found in the supplementary materials.

#### Turning parameter selection

We determine the tuning parameters λ_1_ and λ_2_ (λ_2_ only for CFGL and FGL) according to the Bayesian information criterion (BIC)[66].

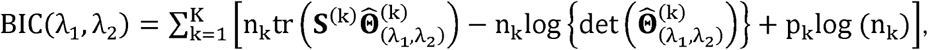

where 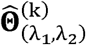 is the estimated precision matrix for the kth condition obtained at (λ_1_,λ_2_), and p_k_ is the number of non-zero elements in 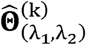. We run the analysis on a series of combinations of λ_1_ and λ_2_, then choose the tuning parameters that achieve the minimal BIC value.

In the simulation, we used BIC to select tuning parameters. For the rat data analysis, because the sample size is very small, it is difficult to obtain meaningful estimates from subsamples in the stability selection. Instead, we first identified the model that achieves the minimum BIC and the models with similar BIC values, then selected the model that have the fewest edges to obtain biologically interpretable results. In the TCGA data analysis, we applied a stability selection procedure to identify reliable edges (see stability selection section).

#### Determining screening matrix

We determined the screening matrix for CFGL by testing the differences between two precision matrix using the method proposed by Xia *et* al.[33]. This method tests whether the difference 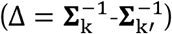 between two precision matrices is 0, i.e.*H*_0_: Δ.0 vs *H*_1_: Δ ≠ 0. To avoid falsely imposing similarity for edges that are moderately differential, we used a relaxed FDR threshold (FDR=0.4) to determine differential entries, such that only the edges that are obviously non-differential across conditions (i.e. FDR>0.4) were encouraged to be similar 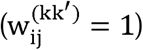. We implemented this method in R and included it in the CFGL package.

#### Generation of Synthetic data

In the simulation study, we generate the gene expression data for multiple network configurations. Suppose each condition contains *M* disjoint modules and each module consists of *p* genes. For each module, the gene constitution is constant across conditions, but the connectivity and the edge weight may vary across conditions. To generate conditions with a specified level of similarity, we first generate the network for condition 1, and then generate the network for other conditions based on their similarities to condition 1.

#### Step 1: Simulating network for condition 1

To generate a module in the network we first create an unweighted scale-free network, according to the Barabasi-Albert model [67] with an exponent of 1, to mimic real-world biological networks structure [68]. Then, we obtain the weighted network 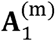 by assigning the edge weights as follows,

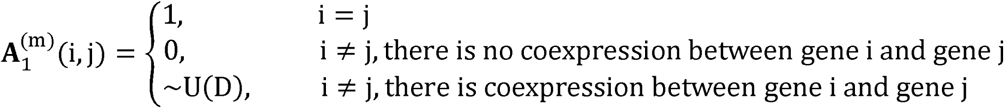

where U(D) is a unifrm distribution with D=[−1, −0.6⋃[0.6,1]. To ensure 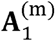 is positive definite, we add values on the matrix diagonal to get the modified matrix 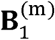:

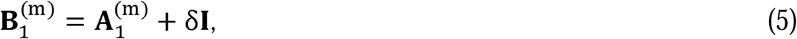

where δ is the minimal eigenvalue of the matrix 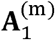. Based on the matrix 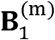, the covariance matrix 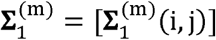 is determined by

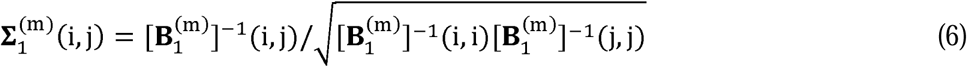

Finally, we obtained the covariance matrix for condition 1, **∑**_1_, by combining 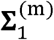 for each module.

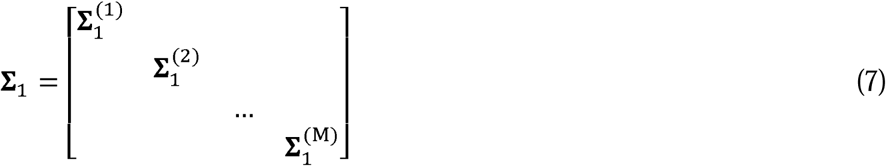

The expression data for condition 1 are generated from N(0,**∑**_1_).

#### Step 2: Simulating network for the other conditions

The modules in other conditions were simulated based on their similarities to the corresponding module in condition 1. Three types of similarities were considered: (1) identical networks structure and identical edge weight across conditions (II), (2) identical network structure but different edge weights across conditions (ID), and (3) different network structures and different edge weights (DD). The generation procedure to obtain 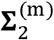 from 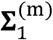 for these three types of similarities is as follows.

1. II modules: the covariance matrix for condition 2 is identical to that for condition 1.

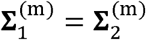
2. ID modules: to maintain the network structure as in condition 1 but altering edge weights, a matrix **U** is added on 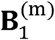:

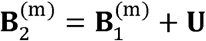

where **U** is a p×p matrix with elements:

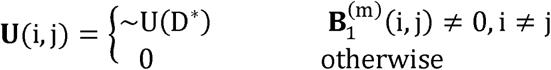

where U(D*) is a uniform distribution with D* = [−0.6,0.6]. Then, 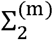 can be obtained from 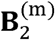 as in (6).
3. DD modules:**∑**_2_ is generated independently as described in step 1.

#### Determining true screening matrix in simulation study

To assess the performance of CFGL-oracle, we obtained the true screening matrix as

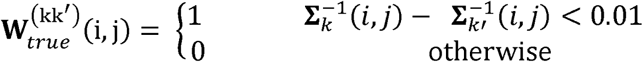

where 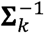 is the simulated precision matrix for the,th condition.

#### Accuracy of edge detection and edge weight estimation in the simulation study

The accuracy of edge identification is assessed by checking if the presence of edges is correct in the estimated in the estimated matrix 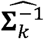. We define true positive as 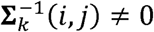 and 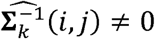 and false positive as 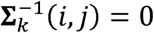 and 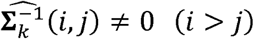. The accuracy of edge weight estimation is assessed by the sum of square error (SSE) between the true and estimated edge weights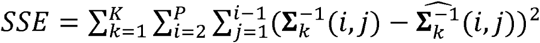 where *K* is number of conditions and *P* is number of nodes (genes). For each *λ*_2_, we generate an ROC curve by computing the true positive rate and false positive rate over a grid of *λ*_1_. Similarly, SSE is computed over a grid of *λ*_1_.

#### Construction of co-expression network for rat data

Heart and brain RNA expression levels were measured in a recombinant inbred (RI) rat panel, HXB/BXH, using the Affymetrix Rat Exon 1.0 ST Array (Affymetrix, Santa Clara, CA). This rat panel was originally generated using gender reciprocal crossing between the congenic Brown Norway strain with the polydactyly-luxate syndrome (BN-Lx/Cub) and the spontaneous hypertensive rat strain (SHR/OlaIpcv), with sixty generations of brother/sister mating after the F2 generation [69]. The CEL files for the heart and brain RNA expression data from 3 to 4 male rats per strain (19 strains) are publically available through the PhenoGen website (http://phenogen.ucdenver.edu) [70] along with a probe mask for the ‘core’ (Affymetrix defined) transcript clusters that eliminates probes that do not align uniquely to the RN6 version of the rat genome or align to a region of the genome that harbors a single nucleotide polymorphism between either of the parental strains (SHR and BN-Lx) and the reference genome. Further detail about this type of probe mask is available in Saba et al 2015 [71]. Transcript cluster estimates on the log base 2 scale were estimated using the rma-sketch pipeline for normalization and aggregation using Affymetrix Power Tools (Irizarry et al 2003; Lockstone 2011) [72][73]. Individual rat estimates were summarized as strain mean values for each transcript cluster and strain combination.

Given the small sample size, we restricted the network construction to the 500 most differentially expressed genes between the two tissues. The differential expression was determined using the R package LIMMA with the default parameter settings.

We ran CFGL and FGL for a grid of *λ*_1_ and *λ*_2_. They both achieved the lowest BIC at *λ*_1_= 0.001 and *λ*_2_ = 0.0008. To investigate the effect of *λ*_2_, we report the results at *λ*_1_= 0.001 and *λ*_2_ = 0.0008,0.0010,0.0012. We set *λ=*.0.0009 for GL since it gives the similar sparsity (edge number) as the other two methods.

#### Construction of co-expression network with TCGA breast cancer data

TCGA has collected gene expression RNA-seq data for 1100 breast cancer patients [43]. We used the normal tissue samples from the individuals (n=112) who have both tumor tissue and matched peripheral normal tissue, and all tumor samples from different individuals that were annotated with ER+ (n=187) and ER-(n=98)[43]. We obtained the gene expression level by downloading the RNA-seq V2 data, which are reads counts normalized by RSEM, from the TCGA website (https://cancergenome.nih.gov/). We then took log transformation for the expression level (with 0.5 added to the counts of each gene to avoid 0) and standardized the transformed expression level to mean 0 and standard deviation 1. Prior to network construction, we first removed genes with very low counts (less or equal than 5) in more than 10% (40) samples. After this step, the log summed read counts over all samples approximately follow a normal distribution.

Due to the limitation of sample size, we restricted our analysis to a subset of 1000 genes. To select 1000 genes in the analysis, we first included 39 genes that were previously reported as breast cancer-related genes (Supplementary Table 6) [23][43]. Then we selected other 961 genes that are most strongly associated with the survival time based on a univariate Cox regression:

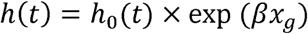

where *t* is the survival time and *x*_g_ is the expression level of the *g*th gene.

#### Stability selection for TCGA data set

In order to obtain reliable co-expression networks, we applied the stability selection procedure in [74] to CFGL, FGL, and GL. This procedure first generates a large set of subsamples from the original data and then builds networks based on the subsamples. The edges that frequently occur in subsamples are kept. This method provides an upper bound for the FDR control and has been shown to outperform the standard GL when being applied to GL [74][37].

In our analysis, we created 100 subsamples, each of which contains half of the original samples. To reduce the computational load, we first determined the optimal choice of *λ*_1_ based on the original dataset, and then used this value for all subsamples. For all methods, *λ*_1 =_ 0.2 achieves both reasonable sparsity and low BIC across a series *λ*_1_ (0.01-0.50) on the original dataset, thus we fixed *λ*_1_ =0.2 in all subsamples. For FGL and CFGL, we performed the analysis on a series of *λ*_2_ for each subsample and then used the tuning parameters that achieve the minimal BIC value to select edges. The minimal BIC for all subsamples were found in the range of *λ*_2_ = 0.002-0.02. We keep the edges that appear in more than 90% subsamples. According to the false discovery rate (FDR) calculation in [37], this threshold guarantees that the number of wrong edges is less than 800 among the 499500 possible edges in the graph.

#### GO enrichment analysis

The GO enrichment analysis were conducted using ToppFun[75], which is publicly available at https://toppgene.cchmc.org/enrichment.jsp. All parameters are used as default.

## Acknowledgement

QL, YL, and FZ are partially supported by the National Institute of Health grant R01GM109453. LX is partially supported by the National Science Foundation grant DMS-1505256. LS is supported by R24AA013162 and P30DA044223. KK is partially supported by R01-AA021131. HK is supported by the National Institute of Health training grant T32 GM102057 awarded to the Pennsylvania State University. YL is also supported by the Huck Graduate Research Innovation Grant from the Pennsylvania State University.

## Supporting information

1. S1_Figures. Supplementary figures
2. S2 _Tables. Supplementary tables
3. S3_ADMM. Detailed ADMM algorithm

## References

1. Blazier AS, Papin JA. Integration of expression data in genome-scale metabolic network reconstructions. Front Physiol. 2012;3:299.

2. Liu L, Lei J, Roeder K. Network assisted analysis to reveal the genetic basis of autism. Ann Appl Stat. 2015;9(3):1571–600.

3. Keller MP, Choi Y, Wang P, Davis DB, Rabaglia ME, Oler AT, et al. A gene expression network model of type 2 diabetes links cell cycle regulation in islets with diabetes susceptibility. Genome Res. 2008;18(5):706–16.

4. Yang Y, Han L, Yuan Y, Li J, Hei N, Liang H. Gene co-expression network analysis reveals common system-level properties of prognostic genes across cancer types. Nat Commun. 2014;5:3231.

5. Pierson E, Koller D, Battle A, Mostafavi S. Sharing and Specificity of Co-expression Networks across 35 Human Tissues. PLoS Comput Biol. 2015;11(5):e1004220.

6. Danaher P, Wang P, Witten DM. The joint graphical lasso for inverse covariance estimation across multiple classes. J R Stat Soc Ser B Stat Method. 2014;76(2):373–97.

7. Xiao X, Moreno-Moral A, Rotival M, Bottolo L, Petretto E. Multi-tissue Analysis of Co-expression Networks by Higher-Order Generalized Singular Value Decomposition Identifies Functionally Coherent Transcriptional Modules. PLoS Genet. 2014;10(1):e1004006.

8. Dobrin R, Zhu J, Molony C, Argman C, Parrish ML, Carlson S, et al. Multi-tissue coexpression networks reveal unexpected subnetworks associated with disease. Genome Biol. 2009;10(5):R55.

9. Li W, Liu C-C, Zhang T, Li H, Waterman MS, Zhou XJ. Integrative Analysis of Many Weighted Co-Expression Networks Using Tensor Computation. PLoS Comput Biol. 2011;7(6):e1001106.

10. Dezso Z, Nikolsky Y, Sviridov E, Shi W, Serebriyskaya T, Dosymbekov D, et al. A comprehensive functional analysis of tissue specificity of human gene expression. BMC Biol. 2008;6(1):49.

11. Messina DN, Glasscock J, Gish W, Lovett M. An ORFeome-based analysis of human transcription factor genes and the construction of a microarray to interrogate their expression. Genome Res. 2004;14(10 B):2041–7.

12. Vaquerizas JM, Kummerfeld SK, Teichmann SA, Luscombe NM. A census of human transcription factors: Function, expression and evolution. Vol. 10, Nature Reviews Genetics. 2009. p. 252–63.

13. Menéndez P, Kourmpetis YAI, ter Braak CJF, van Eeuwijk FA. Gene regulatory networks from multifactorial perturbations using graphical lasso: Application to the DREAM4 challenge. PLoS One. 2010;5(12):e14147.

14. Logsdon BA, Mezey J. Gene expression network reconstruction by convex feature selection when incorporating genetic perturbations. PLoS Comput Biol. 2010;6(12):e1001014.

15. Wang YXR, Huang H. Review on statistical methods for gene network reconstruction using expression data. J Theor Biol. 2014;362:53–61.

16. Fan J, Feng Y, Wu Y. Network exploration via the adaptive LASSO and SCAD penalties. Ann Appl Stat. 2009;3(2):521.

17. Lee KH, Xue L. Nonparametric finite mixture of Gaussian graphical models. Technometrics. 2017;Forthcoming.

18. Ma S, Xue L, Zou H. Alternating direction methods for latent variable Gaussian graphical model selection. Neural Comput. 2013;25(8):2172–98.

19. Xue L, Zou H. Regularized rank-based estimation of high-dimensional nonparanormal graphical models. Ann Stat. 2012;40(5):2541–71.

20. Lauritzen SL. Graphical Models. Clarendon Press; 1996.

21. Langfelder P, Horvath S. WGCNA: an R package for weighted correlation network analysis. BMC Bioinformatics. 2008;9(1):559.

22. Meinshausen N, Bühlmann P. High-dimensional graphs and variable selection with the lasso. Ann Stat. 2006;1436–62.

23. Peng J, Wang P, Zhou N, Zhu J. Partial correlation estimation by joint sparse regression models. J Am Stat Assoc. 2009;104(486):735–46.

24. Guo J, Levina E, Michailidis G, Zhu J. Joint estimation of multiple graphical models. Biometrika. 2011;98(1):1–15.

25. Zhu Y, Shen X, Pan W. Structural pursuit over multiple undirected graphs. J Am Stat Assoc. 2014;109(508):1683–96.

26. Ma J, Michailidis G. Joint structural estimation of multiple graphical models. J Mach Learn Res. 2016;17(166):1–48.

27. Saegusa T, Shojaie A. Joint estimation of precision matrices in heterogeneous populations. Electron J Stat. 2016;10(1):1341.

28. Hoefling H. A path algorithm for the fused lasso signal approximator. J Comput Graph Stat. 2010;19(4):984–1006.

29. Kanehisa M, Goto S. KEGG: kyoto encyclopedia of genes and genomes. Nucleic Acids Res. 2000;28(1):27–30.

30. Okamura Y, Aoki Y, Obayashi T, Tadaka S, Ito S, Narise T, et al. COXPRESdb in 2015: Coexpression database for animal species by DNA-microarray and RNAseq-based expression data with multiple quality assessment systems. Nucleic Acids Res. 2015;43(D1):D82–6.

31. Carlson MR, Zhang B, Fang Z, Mischel PS, Horvath S, Nelson SF. Gene connectivity, function, and sequence conservation: predictions from modular yeast co-expression networks. BMC Genomics. 2006;7:40.

32. Fan J, Lv J. Sure independence screening for ultrahigh dimensional feature space. J R Stat Soc Ser B Stat Method. 2008;70(5):849–911.

33. Xia Y, Cai T, Cai TT. Testing differential networks with applications to the detection of gene-gene interactions. Biometrika. 2015;102(2):247–66.

34. Cai T, Liu W, Luo X. A constrained l 1 minimization approach to sparse precision matrix estimation. J Am Stat Assoc. 2011;106(494):594–607.

35. Friedman J, Hastie T, Tibshirani R. Sparse inverse covariance estimation with the graphical lasso. Biostatistics. 2008;9(3):432–41.

36. Zhao SD, Cai TT, Li H. Direct estimation of differential networks. Biometrika. 2014;101(2):253–68.

37. Meinshausen N, Bühlmann P. Stability selection. J R Stat Soc Ser B Stat Method. 2010;72(4):417–73.

38. Holz A, Schwab ME. Developmental expression of the myelin gene MOBP in the rat nervous system. J Neurocytol. 1997;26(7):467–77.

39. Moss FJ, Dolphin AC, Clare JJ. Human neuronal stargazin-like proteins, γ2, γ3 and γ4; an investigation of their specific localization in human brain and their influence on Ca V 2.1 voltage-dependent calcium channels expressed in Xenopus oocytes. BMC Neurosci. 2003;4(1):23.

40. Everett K V, Chioza B, Aicardi J, Aschauer H, Brouwer O, Callenbach P, et al. Linkage and association analysis of CACNG3 in childhood absence epilepsy. Eur J Hum Genet. 2007;15(4):463–72.

41. Tamura N, Ogawa Y, Chusho H, Nakamura K, Nakao K, Suda M, et al. Cardiac fibrosis in mice lacking brain natriuretic peptide. Proc Natl Acad Sci. 2000;97(8):4239–44.

42. Wang Q, Lin JL-C, Chan SY, Lin JJ-C. The Xin repeat-containing protein, mXinβ, initiates the maturation of the intercalated discs during postnatal heart development. Dev Biol. 2013;374(2):264–80.

43. The Cancer Genome Atlas Network. Comprehensive molecular portraits of human breast tumors. Nature. 2012;490(7418):61–70.

44. Gruvberger S, Ringner M, Chen Y, Panavally S, Saal LH, Borg A, et al. Estrogen receptor status in breast cancer is associated with remarkably distinct gene expression patterns. Cancer Res. 2001;61:5979–84.

45. Zhang Z, Christin JR, Wang C, Ge K, Oktay MH, Guo W. Mammary-Stem-Cell-Based Somatic Mouse Models Reveal Breast Cancer Drivers Causing Cell Fate Dysregulation. Cell Rep. 2016;16(12):3146–56.

46. Eelen G, Vanden Bempt I, Verlinden L, Drijkoningen M, Smeets A, Neven P, et al. Expression of the BRCA1-interacting protein Brip1/BACH1/FANCJ is driven by E2F and correlates with human breast cancer malignancy. Oncogene. 2008;27(30):4233–41.

47. Zhang Y, Zang M, Li J, Ji J, Zhang J, Liu X, et al. CEACAM6 promotes tumor migration, invasion, and metastasis in gastric cancer. Acta Biochim Biophys Sin. 2014;46(4):283–90.

48. Vecchi M, Confalonieri S, Nuciforo P, Vigano MA, Capra M, Bianchi M, et al. Breast cancer metastases are molecularly distinct from their primary tumors. Oncogene. 2008;27(15):2148–58.

49. Bezault J, Bhimani R, Wiprovnick J, Furmanski P. Human lactoferrin inhibits growth of solid tumors and development of experimental metastases in mice. Cancer Res. 1994;54(9):2310–2.

50. Ushida Y, Sekine K, Kuhara T, Takasuka N, Iigo M, Tsuda H. Inhibitory effects of bovine lactoferrin on intestinal polyposis in the Apc Min mouse. Cancer Lett. 1998;134(2):141–5.

51. Jung HC, Kim SH, Lee JH, Kim JH, Han SW. Gene Regulatory Network Analysis for Triple-Negative Breast Neoplasms by Using Gene Expression Data. J Breast Cancer. 2017;20(3):240–5.

52. Moiola C, De Luca P, Gardner K, Vazquez E, De Siervi A. Cyclin T1 overexpression induces malignant transformation and tumor growth. Cell Cycle. 2010;9(15):3191–8.

53. Xiong D, Li G, Li K, Xu Q, Pan Z, Ding F, et al. Exome sequencing identifies MXRA5 as a novel cancer gene frequently mutated in non--small cell lung carcinoma from Chinese patients. Carcinogenesis. 2012;33(9):1797–805.

54. Wang G, Yao L, Xu H, Tang W, Fu J, Hu X, et al. Identification of MXRA5 as a novel biomarker in colorectal cancer. Oncol Lett. 2013;5(2):544–8.

55. Cantor SB, Guillemette S. Hereditary breast cancer and the BRCA1-associated FANCJ/BACH1/BRIP1. Futur Oncol. 2011;7(2):253–61.

56. Liang Y, Wu H, Lei R, Chong RA, Wei Y, Lu X, et al. Transcriptional network analysis identifies BACH1 as a master regulator of breast cancer bone metastasis. J Biol Chem. 2012;287(40):33533–44.

57. Yao Z, Luo J, Hu K, Lin J, Huang H, Wang Q, et al. ZKSCAN1 gene and its related circular RNA (circ ZKSCAN1) both inhibit hepatocellular carcinoma cell growth, migration, and invasion but through different signaling pathways. Mol Oncol. 2017;11(4):422–37.

58. Fan L, Tan B, Li Y, Zhao Q, Liu Y, Wang D, et al. Silencing of ZNF139-siRNA induces apoptosis in human gastric cancer cell line BGC823. Int J Clin Exp Pathol. 2015;8(10):12428–36.

59. Blumenthal RD, Hansen HJ, Goldenberg DM. Inhibition of adhesion, invasion, and metastasis by antibodies targeting CEACAM6 (NCA-90) and CEACAM5 (Carcinoembryonic Antigen). Cancer Res. 2005;65(19):8809–17.

60. Duxbury MS, Matros E, Clancy T, Bailey G, Doff M, Zinner MJ, et al. CEACAM6 Is a Novel Biomarker in Pancreatic Adenocarcinoma and PanIN Lesions. Ann Surg. 2005;241(3):491–6.

61. Kohno D, Yada T. Arcuate NPY neurons sense and integrate peripheral metabolic signals to control feeding. Neuropeptides. 2012;46(6):315–9.

62. Liu L, Xu Q, Cheng L, Ma C, Xiao L, Xu D, et al. NPY1R is a novel peripheral blood marker predictive of metastasis and prognosis in breast cancer patients. Oncol Lett. 2015;9(2):891–6.

63. Höfling H, Tibshirani R. Estimation of sparse binary pairwise markov networks using pseudo-likelihoods. J Mach Learn Res. 2009;10(Apr):883–906.

64. Xue L, Zou H, Cai T, others. Nonconcave penalized composite conditional likelihood estimation of sparse Ising models. Ann Stat. 2012;40(3):1403–29.

65. Boyd S, Parikh N, Chu E, Peleato B, Eckstein J. Distributed optimization and statistical learning via the alternating direction method of multipliers. Found Trends Mach Learn. 2011;3(1):1–122.

66. Schwarz G, others. Estimating the dimension of a model. Ann Stat. 1978;6(2):461–4.

67. Albert R, Barabási A-L. Statistical mechanics of complex networks. Rev Mod Phys. 2002;74(1):47.

68. Newman MEJ. The structure and function of complex networks. SIAM Rev. 2003;45(2):167–256.

69. Printz MP, Jirout M, Jaworski R, Alemayehu A, Kren V. Invited Review: HXB/BXH rat recombinant inbred strain platform: a newly enhanced tool for cardiovascular, behavioral, and developmental genetics and genomics. J Appl Physiol. 2003;94(6):2510–22.

70. Hoffman PL, Bennett B, Saba LM, Bhave S V, Carosone-Link PJ, Hornbaker CK, et al. Using the Phenogen website for “in silico”analysis of morphine-induced analgesia: identifying candidate genes. Addict Biol. 2011;16(3):393–404.

71. Saba LM, Flink SC, Vanderlinden LA, Israel Y, Tampier L, Colombo G, et al. The sequenced rat brain transcriptome--its use in identifying networks predisposing alcohol consumption. FEBS J. 2015;282(18):3556–78.

72. Irizarry RA, Bolstad BM, Collin F, Cope LM, Hobbs B, Speed TP. Summaries of Affymetrix GeneChip probe level data. Nucleic Acids Res. 2003;31(4):e15.

73. Lockstone HE. Exon array data analysis using Affymetrix power tools and R statistical software. Brief Bioinform. 2011;12(6):634–44.

74. Shah RD, Samworth RJ. Variable selection with error control: another look at stability selection. J R Stat Soc Ser B (Stat Method). 2013;75(1):55–80.

75. Chen J, Bardes EE, Aronow BJ, Jegga AG. ToppGene Suite for gene list enrichment analysis and candidate gene prioritization. Nucleic Acids Res. 2009;37(suppl_2):W305-W311.

