## Supplementary Materials for "Condition-adaptive fused graphical lasso (CFGL): an adaptive procedure for inferring condition-specific gene co-expression network"

**Supplementary Figures**

**Figure 1. Comparison of performance for simulations with two conditions with sample size n=100. Top row: ROC curves for edge detection. Bottom row: SSE for edge weight estimation. Red line: CFGL, Green line: FGL, Blue line: GL, Purple line: CFGL-oracle.**

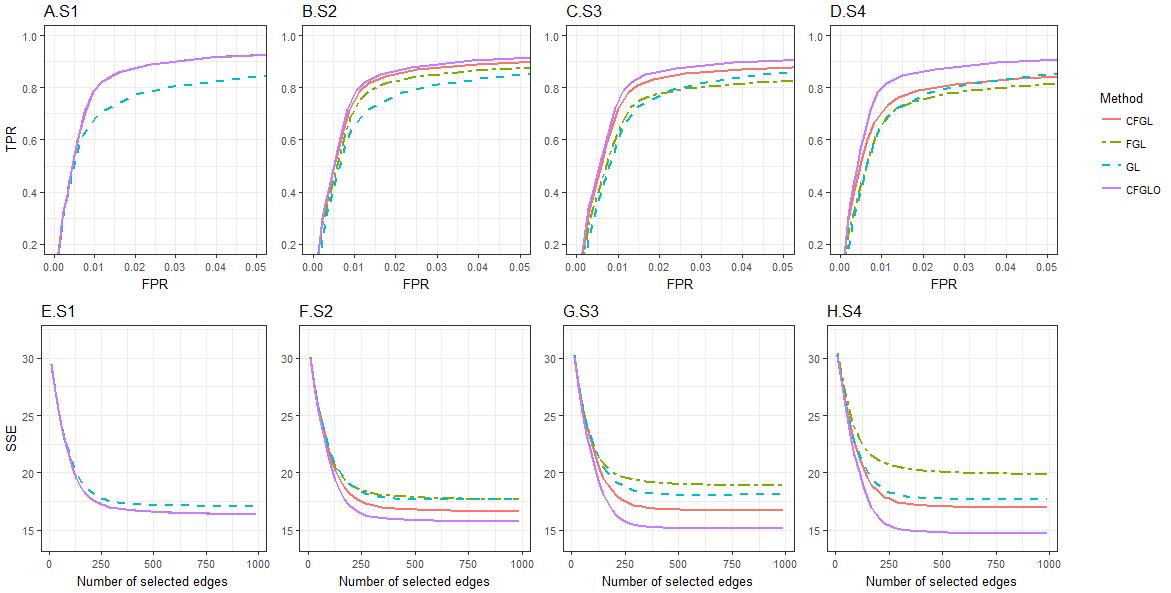

**Figure 2. The rat brain specific network.**

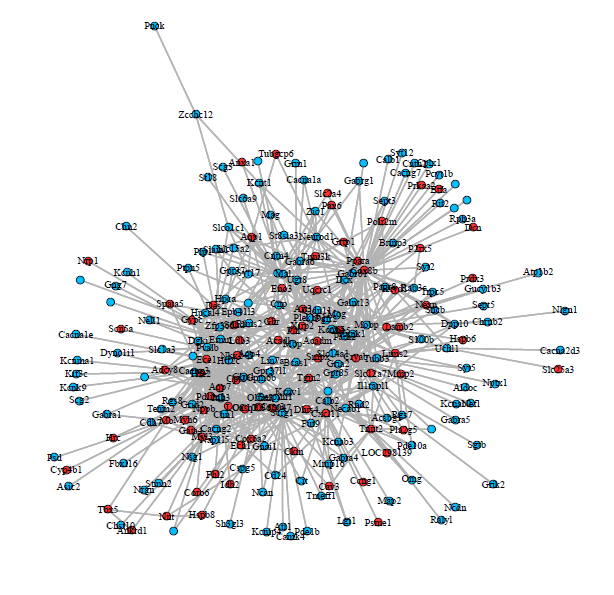

**Figure 3. The rat heart specific network.**

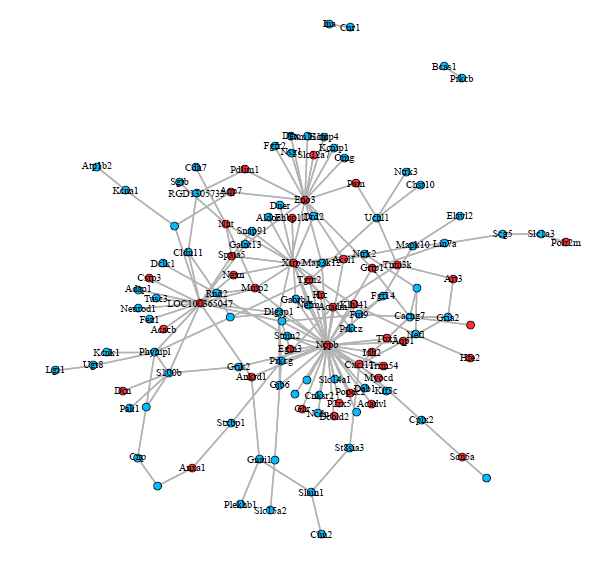

**Figure 4. The ER+/ER- shared subnetwork. Red: Genes that are up-regulated in both tumor tissues in comparison with normal tissue. Blue: Genes that are down-regulated in both tumor tissues. Yellow: Genes that are up-regulated in one tumor tissue but down-regulated in another.**

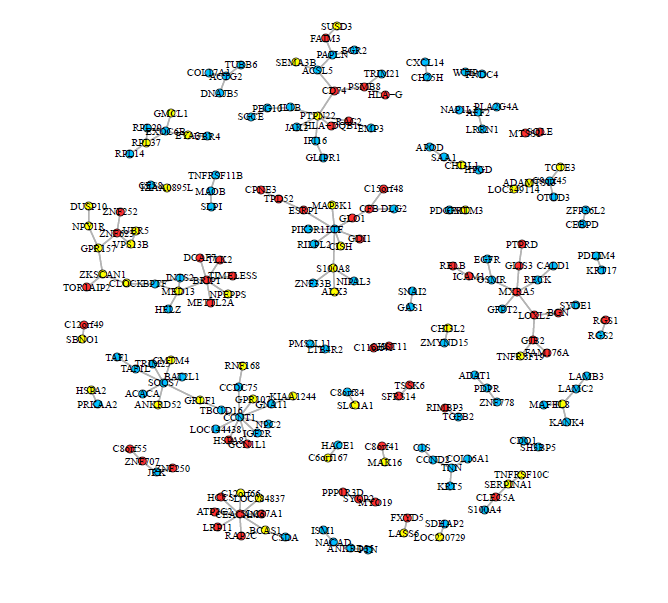

**Figure 5. The ER+ specific subnetwork. Red: Genes that are up-regulated in both tumor tissues in comparison with normal tissue. Blue: Genes that are down-regulated in both tumor tissues. Yellow: Genes that are up-regulated in one tumor tissue but down-regulated in another.**

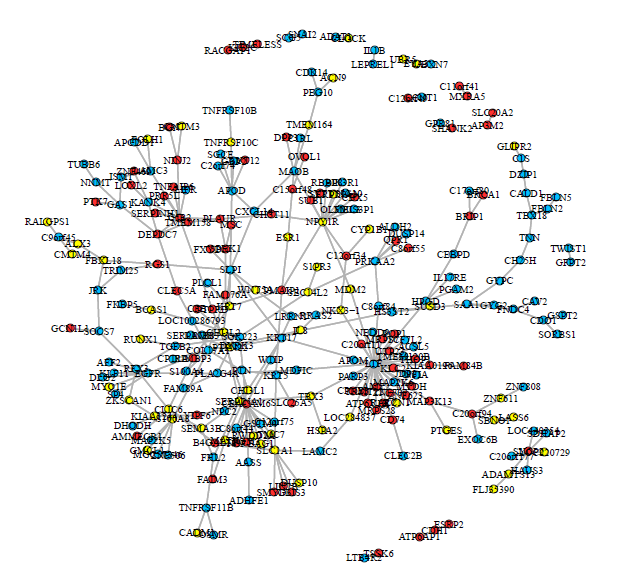

**Figure 6. The ER- specific subnetwork. Red: Genes that are up-regulated in both tumor tissues in comparison with normal tissue. Blue: Genes that are down-regulated in both tumor tissues. Yellow: Genes that are up-regulated in one tumor tissue but down-regulated in another.**

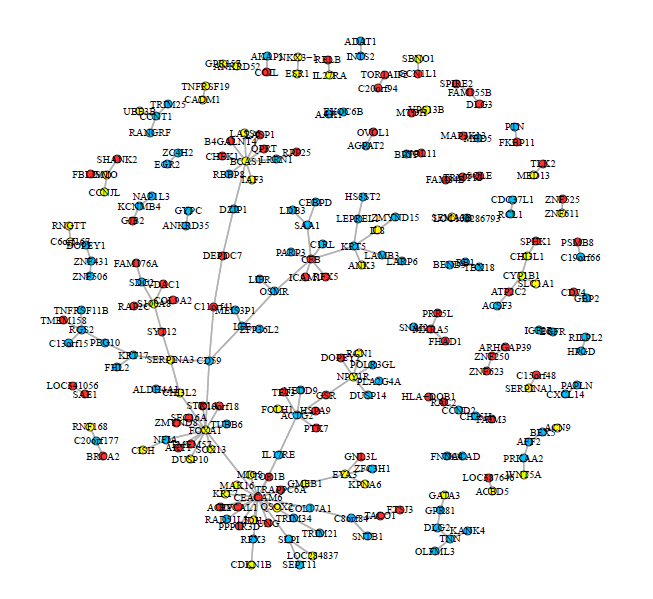
