## Supplementary Materials for "Condition-adaptive fused graphical lasso (CFGL): an adaptive procedure for inferring condition-specific gene co-expression network"

**Supplementary Tables**

**Table 1. Comparison of partial AUCs in the 2-condition simulation studies.**

CFGL, CFGLO, FGL and GL were compared under 4 simulation scenarios (S1-S4) with 2 sample sizes (n=50 and 100). The analysis was ran on a grid of $\lambda_{1}$ and $\lambda_{2}$. The ROC curves were computed over $\lambda_{1}$ for each fixed $\lambda_{2}$, and the partial AUCs in the FPR range of [0, 0.05] were computed. The table summarizes the partial AUCs for $\lambda_{2}=$0.05, 0.10, 0.15, and 0.20.

| $\lambda_{2}$ | Method | n = 50 | | | | n = 100 | | | |
| --- | --- | --- | --- | --- | --- | --- | --- | --- | --- |
|  |  | S1 | S2 | S3 | S4 | S1 | S2 | S3 | S4 |
| 0.05 | CFGL | 0.650 | 0.641 | 0.644 | 0.619 | 0.786 | 0.748 | 0.765 | 0.740 |
|  | CFGLO | 0.651 | 0.656 | 0.652 | 0.658 | 0.786 | 0.760 | 0.771 | 0.775 |
|  | FGL | 0.651 | 0.619 | 0.634 | 0.606 | 0.786 | 0.719 | 0.752 | 0.721 |
|  | GL | 0.583 | 0.594 | 0.588 | 0.590 | 0.726 | 0.702 | 0.713 | 0.714 |
| 0.10 | CFGL | 0.692 | 0.663 | 0.676 | 0.626 | 0.811 | 0.762 | 0.784 | 0.739 |
|  | CFGLO | 0.694 | 0.695 | 0.693 | 0.711 | 0.811 | 0.787 | 0.798 | 0.806 |
|  | FGL | 0.694 | 0.621 | 0.656 | 0.600 | 0.811 | 0.704 | 0.756 | 0.700 |
|  | GL | 0.583 | 0.594 | 0.588 | 0.590 | 0.726 | 0.702 | 0.713 | 0.714 |
| 0.15 | CFGL | 0.711 | 0.671 | 0.688 | 0.620 | 0.817 | 0.764 | 0.788 | 0.730 |
|  | CFGLO | 0.714 | 0.717 | 0.715 | 0.723 | 0.817 | 0.800 | 0.808 | 0.820 |
|  | FGL | 0.714 | 0.609 | 0.660 | 0.578 | 0.817 | 0.677 | 0.747 | 0.665 |
|  | GL | 0.583 | 0.594 | 0.588 | 0.590 | 0.726 | 0.702 | 0.713 | 0.714 |
| 0.20 | CFGL | 0.718 | 0.671 | 0.691 | 0.612 | 0.818 | 0.810 | 0.789 | 0.722 |
|  | CFGLO | 0.721 | 0.729 | 0.724 | 0.735 | 0.818 | 0.765 | 0.814 | 0.828 |
|  | FGL | 0.721 | 0.589 | 0.653 | 0.550 | 0.818 | 0.649 | 0.735 | 0.631 |
|  | GL | 0.583 | 0.594 | 0.588 | 0.590 | 0.726 | 0.702 | 0.713 | 0.714 |

**Table 2. Comparison of partial AUCs in the 3-condition simulation studies.**

CFGL, FGL and GL were compared under 4 simulation scenarios (S1-S4) with n=50. The analysis was ran on a grid of $\lambda_{1}$ and $\lambda_{2}$. The minimum BIC was achieved at $\lambda_{2}$=0.15. The ROC curves were computed over $\lambda_{1}$ with $\lambda_{2}$=0.15. The table summarized the partial AUCs in the FPR range of [0, 0.05].

| Method | S1 | S2 | S3 | S4 |
| --- | --- | --- | --- | --- |
| CFGL | 0.649 | 0.508 | 0.650 | 0.513 |
| FGL | 0.605 | 0.494 | 0.615 | 0.499 |
| GL | 0.597 | 0.474 | 0.593 | 0.472 |

**Table 3. Numbers of detected co-expression edges in the rat multi-tissue dataset.**

The optimal BIC was achieved at $\lambda_{1}=$0.0010 and $\lambda_{2}=$0.0008 for CFGL and FGL, and $\lambda_{1}=$0.0009 for GL. We investigated the effect of $\lambda_{2}$by repeating the analysis for CFGL and FGL at $\lambda_{2}=$0.0010 and 0.0012.

| Method | $\lambda_{1}$ | $\lambda_{2}$ | Brain specific | Heart specific | Common |
| --- | --- | --- | --- | --- | --- |
| CFGL | 0.0010 | 0.0008 | 815 | 203 | 356 |
| FGL |  |  | 611 | 29 | 354 |
| CFGL | 0.0010 | 0.0010 | 446 | 293 | 522 |
| FGL |  |  | 233 | 15 | 522 |
| CFGL | 0.0010 | 0.0012 | 280 | 188 | 617 |
| FGL |  |  | 57 | 7 | 623 |
| GL | 0.0009 | - | 883 | 605 | 3 |

**Table 4. Top-5 tissue-specific hubs identified by GL with rat expression data.**

| Tissue | Hubs | CFGL | | FGL | | GL | |
| --- | --- | --- | --- | --- | --- | --- | --- |
|  |  | #edge | #edge  ranking | #edge | #edge  Ranking | #edge | #edge  Ranking |
| Brain | Fh12 | 4 | 90 | 3 | 92 | 67 | 1 |
|  | Neurod1 | 15 | 28 | 13 | 29 | 60 | 2 |
|  | Camkv | 0 | - | 0 | - | 34 | 3 |
|  | Elavl3 | 0 | - | 0 | - | 30 | 4 |
|  | Xirp2 | 4 | 92 | 0 | - | 27 | 5 |
| Heart | Cacna2d3 | 0 | - | 0 | - | 41 | 1 |
|  | Ckm | 0 | - | 0 | - | 35 | 2 |
|  | Olfm1 | 0 | - | 0 | - | 32 | 3 |
|  | Rit2 | 0 | - | 0 | - | 31 | 4 |
|  | Camkk2 | 0 | - | 0 | - | 30 | 5 |

**Table 5. Disease type specificity of the estimated co-expression edges for the TCGA breast cancer data.**

|  | Normal only | ER+ tumor only | ER- tumor only | Shared in normal and ER+ | Shared in normal and ER- | Shared in ER+ and ER- | Shared in all tissue | Total number of edges |
| --- | --- | --- | --- | --- | --- | --- | --- | --- |
| CFGL | 384 | 554 | 360 | 250 | 60 | 332 | 684 | 2624 |
| FGL | 1034 | 840 | 472 | 446 | 100 | 226 | 1330 | 4448 |
| GL | 2250 | 2136 | 980 | 290 | 36 | 218 | 68 | 5978 |

**Table 6. List of the 39 genes that are known to be related to breast cancer and are included in our TCGA analysis**

| Gene | Source |  | Gene | Source |  | Gene | Source |
| --- | --- | --- | --- | --- | --- | --- | --- |
| FOXA1 | 1 |  | CDKN1B | 2 |  | BRCA2 | 2 |
| KIF2C | 1 |  | MAP2K4 | 2 |  | BRIP1 | 2 |
| AURKB | 1 |  | TBX3 | 2 |  | CHEK2 | 2 |
| RAD54L | 1 |  | CBFB | 2 |  | NBN | 2 |
| BUB1 | 1 |  | AFF2 | 2 |  | AKT1 | 2 |
| PTEN | 2 |  | PIK3R1 | 2 |  | AKT2 | 2 |
| AKT1 | 2 |  | PTPN22 | 2 |  | ESR1 | 2 |
| TP53 | 2 |  | PTPRD | 2 |  | MDM2 | 2 |
| GATA3 | 2 |  | NF1 | 2 |  | TUBB1 | 2 |
| CDH1 | 2 |  | SF3B1 | 2 |  | JAK2 | 2 |
| RB1 | 2 |  | CCND3 | 2 |  | EGFR | 2 |
| MLL3 | 2 |  | ATM | 2 |  | MAP2K1 | 2 |
| MAP3K1 | 2 |  | BRCA1 | 2 |  | CHEK1 | 2 |

Source 1: Peng, J., Wang, P., Zhou, N., & Zhu, J. (2009). Partial correlation estimation by joint sparse regression models. Journal of the American Statistical Association, 104(486), 735-746.

Source 2: Cancer Genome Atlas Network. (2012). Comprehensive molecular portraits of human breast tumors. Nature, 490(7418), 61
