## Supplementary Materials for "Condition-adaptive fused graphical lasso (CFGL): an adaptive procedure for inferring condition-specific gene co-expression network"

### Supporting Infomation 3

#### Appendix A ADMM Algorithm For CFGL

In this section, we present the details of algorithm to solve the problem (2)

$$\max_{\{\Theta\}}\left(\sum_{k=1}^{K} n_k\left[\log\left(\det(\boldsymbol{\Theta}^{(k)})-\operatorname{tr}(\boldsymbol{S}^{(k)}\boldsymbol{\Theta}^{(k)})\right)\right]-P\left\{\boldsymbol{\Theta}\right\}\right),$$

where the penalty term is the fused lasso

$$P\left\{\boldsymbol{\Theta}\right\}=\lambda_1\sum_{k=1}^{K}\sum_{i\neq j}|\theta_{ij}^{(k)}|+\lambda_2\sum_{k<k'}^{K}\sum_{i\neq j} w_{ij}^{(kk')}|\theta_{ij}^{(k)}-\theta_{ij}^{(k')}|.$$

Following Danaher et al. (2014), we solve problem (2) by using ADMM algorithm (Boyd et al., 2010). To proceed, we rewrite the problem as

$$\min_{\{\Theta\},\{Z\}}\left(-\sum_{k=1}^{K} n_k\left[\log\left(\det(\boldsymbol{\Theta}^{(k)})-\operatorname{tr}\left(\boldsymbol{S}^{(k)}\boldsymbol{\Theta}^{(k)}\right)\right)\right]+P\{\boldsymbol{Z}\}\right)$$

subject to the positive definiteness constraint and $\boldsymbol{Z}^{(k)}=\boldsymbol{\Theta}^{k}$ for $k=1,\ldots,K$, where $\{\boldsymbol{Z}\}=\left(\boldsymbol{Z}^{(1)},\ldots,\boldsymbol{Z}^{(K)}\right)$. By Boyd et al. (2010), the scaled augmented Lagrangian for this problem is

$$\begin{aligned}L\left(\{\boldsymbol{\Theta}\},\{\boldsymbol{Z}\},\{\boldsymbol{U}\}\right)=&-\sum_{k=1}^{K} n_k\left[\log\left(\det(\boldsymbol{\Theta}^{(k)})-\operatorname{tr}\left(\boldsymbol{S}^{(k)}\boldsymbol{\Theta}^{(k)}\right)\right)\right]+P\{\boldsymbol{Z}\}\\&+\frac{\rho}{2}\sum_{k=1}^{K}\|\boldsymbol{\Theta}^{(k)}-\boldsymbol{Z}^{(k)}+\boldsymbol{U}^{(k)}\|_F^2-\frac{\rho}{2}\sum_{k=1}^{K}\|\boldsymbol{U}^{(k)}\|_F^2,\end{aligned}$$

where $\{\boldsymbol{U}\}=\boldsymbol{U}^{(1)},\ldots,\boldsymbol{U}^{(k)}$ are dual variables and $\rho$ serves as a 'penalty parameter', and $\|\boldsymbol{\Theta}\|$ is the Frobenius norm of matrix $\boldsymbol{\Theta}$, $\|\boldsymbol{\Theta}\|_F^2=\sum_i\sum_j\Theta_{ij}^2$.

Following the procedure of ADMM algorithm for JGL problem (Danaher et al., 2014), we present the ADMM algorithm for solving problem (2).

Step 1 Initialize values: $\boldsymbol{\Theta}^{(k)}=\boldsymbol{I},\boldsymbol{U}^{(k)}=\boldsymbol{0}$ and $\boldsymbol{Z}^{(k)}=\boldsymbol{0}$ for $k=1,\ldots,K$. Set $\rho>0$.

Step 2 For the $i$th step, update the following procedures.

(i) For $k = 1, \dots, K$, update $\mathbf{\Theta}_{(i)}^{(k)}$ as the minimizete of

$$-n_k \left[\log(\det(\mathbf{\Theta}^{(k)})) - \mathrm{tr}(\boldsymbol{S}^{(k)}\mathbf{\Theta}^{(k)})\right] + \frac{\rho}{2}\|\mathbf{\Theta}^{(k)} - \boldsymbol{Z}_{(i-1)}^{(k)} + \boldsymbol{U}_{(i-1)}^{(k)}\|_F^2.$$

Let $\boldsymbol{V}\boldsymbol{D}\boldsymbol{V}^\top$ be the eigendecomposition of $\boldsymbol{S}^{(k)} - \rho\boldsymbol{Z}_{(i-1)}^{k}/n_k + \rho\boldsymbol{U}_{(i-1)}^{(k)}/n_k$, the solution is given by (Witten and Tibshirani, 2009) by $\boldsymbol{V}\widetilde{\boldsymbol{D}}\boldsymbol{V}^\top$, where $\widetilde{\boldsymbol{D}}$ is the diagonal matrix with $j$th diagonal element $\frac{n_k}{2\rho}\left[-D_{jj} + \sqrt{D_{jj}^2 + 4\rho/n_k}\right]$.

(ii) Update $\{\boldsymbol{Z}_{(i)}\}$ as the minimizer of

$$\frac{\rho}{2}\sum_{k=1}^{K}\|\boldsymbol{Z}^{(k)} - (\mathbf{\Theta}_{(i)}^{(k)} + \boldsymbol{U}_{(i-1)}^{(k)})\|_F^2 + P(\{\boldsymbol{Z}\}). \tag{A.1}$$

(iii) For $k = 1, \dots, K$, update $\boldsymbol{U}_{(i)}^{(k)} = \boldsymbol{U}_{(i-1)}^{(k)} + \mathbf{\Theta}_{(i)}^{(k)} - \boldsymbol{Z}_{(i)}^{(k)}$.

Step 3 Repeat Step 2 until convergence.

*Remarks.*

(1) This algorithm is guaranteed to converge to the global optimum, by Boyd et al. (2010). The positive definiteness constraint on the estimated precision matrices is naturally enforced by Step 2 (i).

(2) We use $\rho = 1$ for controlling the step size and declare convergence when

$$\sum_k \|\mathbf{\Theta}_{(i)}^{(k)} - \mathbf{\Theta}_{(i-1)}^{(k-1)}\|_1 / \sum_k \|\mathbf{\Theta}_{(i-1)}^{(k-1)}\|_1 < 10^{-5}.$$

(3) It remains the task of minimizing (A.1). Note that the problem (A.1) can be rewritten as

$$\min_{\{Z\}}\left[\frac{\rho}{2}\sum_{k=1}^{K}\|\boldsymbol{Z}^{(k)} - \boldsymbol{A}^{(k)}\|_F^2 + \lambda_1\sum_{k=1}^{K}\sum_{i\neq j}|Z_{ij}^{(k)}| + \lambda_2\sum_{k<k'}\sum_{i\neq j}w_{ij}^{(kk')}|Z_{ij}^{(k)} - Z_{ij}^{(k')}|\right], \tag{A.2}$$

where $\boldsymbol{A}^{(k)} = \mathbf{\Theta}_{(i)}^{(k)} + \boldsymbol{U}_{(i-1)}^{(k)}$.

Since the problem (A.2) is completely separable with respect to each pair of matrix element $(i, j)$, one can simply solve the problem

$$\min_{Z_{ij}^{(1)},\dots,Z_{ij}^{(K)}}\left[\frac{\rho}{2}\sum_{k=1}^{K}(\boldsymbol{Z}_{ij}^{(k)} - \boldsymbol{A}_{ij}^{(k)}\|_F^2 + \lambda_1\sum_{k=1}^{K}1_{i\neq j}|Z_{ij}^{(k)}| + \lambda_2\sum_{k<k'}w_{ij}^{(kk')}|Z_{ij}^{(k)} - Z_{ij}^{(k')}|\right], \tag{A.3}$$

for each $(i, j)$.

Let $\widehat{\boldsymbol{Z}}(\lambda_1, \lambda_2) = \left(\widehat{Z}_{ij}^{(1)}(\lambda_1, \lambda_2), \dots, \widehat{Z}_{ij}^{(K)}(\lambda_1, \lambda_2)\right)^\top$ be the solution to the problem (A.3). Note that $\widehat{\boldsymbol{Z}}(\lambda_1, \lambda_2)$ depends on $\lambda_1$ and $\lambda_2$. By Friedman et al. (2007), $\widehat{\boldsymbol{Z}}(\lambda_1, \lambda_2) = \mathrm{SH}\left(\widehat{\boldsymbol{Z}}(0, \lambda_2), \frac{\lambda_1}{\rho}\right)$, where the soft thresholding operator $\text{SH}(x, \lambda)$ is defined by

$$\text{SH}(x, \lambda) = \begin{cases} x + \lambda, & x < \lambda, \\ 0, & -\lambda < x < \lambda, \\ x - \lambda, & x > \lambda. \end{cases}$$

We present two examples for illustration to derive $\widehat{\boldsymbol{Z}}(0, \lambda_2)$ for $K = 2$ and $K = 3$, respectively.

**Example 1**. For $K = 2$, the solution $\widehat{\boldsymbol{Z}}(0, \lambda_2) = \left(\widehat{Z}_{ij}^{(1)}, \widehat{Z}_{ij}^{(2)}\right)$ takes the form

(1) If $A_{ij}^{(1)} - A_{ij}^{(2)} > \frac{2\lambda_2}{\rho} w_{ij}^{(12)}$, then

$$\left(\widehat{Z}_{ij}^{(1)}, \widehat{Z}_{ij}^{(2)}\right) = \left(A_{ij}^{(1)} - \frac{\lambda_2}{\rho} w_{ij}^{(12)}, A_{ij}^{(1)} + \frac{\lambda_2}{\rho} w_{ij}^{(12)}\right);$$

(2) If $A_{ij}^{(1)} - A_{ij}^{(2)} < -\frac{2\lambda_2}{\rho} w_{ij}^{(12)}$, then

$$\left(\widehat{Z}_{ij}^{(1)}, \widehat{Z}_{ij}^{(2)}\right) = \left(A_{ij}^{(1)} + \frac{\lambda_2}{\rho} w_{ij}^{(12)}, A_{ij}^{(1)} - \frac{\lambda_2}{\rho} w_{ij}^{(12)}\right);$$

(3) If $|A_{ij}^{(1)} - A_{ij}^{(2)}| \leq \frac{2\lambda_2}{\rho} w_{ij}^{(12)}$, then

$$\left(\widehat{Z}_{ij}^{(1)}, \widehat{Z}_{ij}^{(2)}\right) = \left(\frac{A_{ij}^{(1)} + A_{ij}^{(2)}}{2}, \frac{A_{ij}^{(1)} + A_{ij}^{(2)}}{2}\right).$$

**Example 2**. For $K = 3$, the solution $\widehat{\boldsymbol{Z}}(0, \lambda_2) = \left(\widehat{Z}_{ij}^{(1)}, \widehat{Z}_{ij}^{(2)}, \widehat{Z}_{ij}^{(3)}\right)$ takes the form

(1) If $A_{ij}^{(1)} - \frac{\lambda_2}{\rho} w_{ij}^{(12)} - \frac{\lambda_2}{\rho} w_{ij}^{(13)} > A_{ij}^{(2)} + \frac{\lambda_2}{\rho} w_{ij}^{(12)} - \frac{\lambda_2}{\rho} w_{ij}^{(23)} > A_{ij}^{(3)} + \frac{\lambda_2}{\rho} w_{ij}^{(23)} + \frac{\lambda_2}{\rho} w_{ij}^{(13)}$, then

$$\left(\widehat{Z}_{ij}^{(1)}, \widehat{Z}_{ij}^{(2)}, \widehat{Z}_{ij}^{(3)}\right) = \left(A_{ij}^{(1)} - \frac{\lambda_2}{\rho} w_{ij}^{(12)} - \frac{\lambda_2}{\rho} w_{ij}^{(13)}, A_{ij}^{(2)} + \frac{\lambda_2}{\rho} w_{ij}^{(12)} - \frac{\lambda_2}{\rho} w_{ij}^{(23)}, A_{ij}^{(3)} + \frac{\lambda_2}{\rho} w_{ij}^{(23)} + \frac{\lambda_2}{\rho} w_{ij}^{(13)}\right);$$

(2) If $|A_{ij}^{(1)} - A_{ij}^{(2)} - \frac{\lambda_2}{\rho} w_{ij}^{(13)} + \frac{\lambda_2}{\rho} w_{ij}^{(23)}| \leq \frac{\lambda_2}{\rho} w_{ij}^{(12)}$, and

$$\frac{A_{ij}^{(1)} + A_{ij}^{(2)} - \frac{\lambda_2}{\rho}(w_{ij}^{(13)} + w_{ij}^{(23)})}{2} > A_{ij}^{(3)} + \frac{\lambda_2}{\rho}(w_{ij}^{(23)} + w_{ij}^{(13)}),$$

then

$$\begin{aligned} &\left(\widehat{Z}_{ij}^{(1)}, \widehat{Z}_{ij}^{(2)}, \widehat{Z}_{ij}^{(3)}\right) \\ = &\left(\frac{A_{ij}^{(1)} + A_{ij}^{(2)} - \frac{\lambda_2}{\rho}(w_{ij}^{(13)} + w_{ij}^{(23)})}{2}, \frac{A_{ij}^{(1)} + A_{ij}^{(2)} - \frac{\lambda_2}{\rho}(w_{ij}^{(13)} + w_{ij}^{(23)})}{2}, A_{ij}^{(3)} + \frac{\lambda_2}{\rho}(w_{ij}^{(23)} + w_{ij}^{(13)})\right); \end{aligned}$$

(3) If $|A_{ij}^{(2)} - A_{ij}^{(3)} + \frac{\lambda_2}{\rho} w_{ij}^{(12)} - \frac{\lambda_2}{\rho} w_{ij}^{(13)}| \leq \frac{\lambda_2}{\rho} w_{ij}^{(23)}$, and

$$\frac{A_{ij}^{(2)} + A_{ij}^{(3)} + \frac{\lambda_2}{\rho}(w_{ij}^{(12)} + w_{ij}^{(13)})}{2} < A_{ij}^{(1)} - \frac{\lambda_2}{\rho}(w_{ij}^{(12)} + w_{ij}^{(13)}),$$

then

$$\begin{aligned} & \left(\widehat{Z}_{ij}^{(1)}, \widehat{Z}_{ij}^{(2)}, \widehat{Z}_{ij}^{(3)}\right) \\ = & \left(A_{ij}^{(1)} - \frac{\lambda_2}{\rho}(w_{ij}^{(12)} + w_{ij}^{(13)}), \frac{A_{ij}^{(2)} + A_{ij}^{(3)} - \frac{\lambda_2}{\rho}(w_{ij}^{(12)} + w_{ij}^{(13)})}{2}, \frac{A_{ij}^{(2)} + A_{ij}^{(3)} - \frac{\lambda_2}{\rho}(w_{ij}^{(12)} + w_{ij}^{(13)})}{2}\right); \end{aligned}$$

(4) If $A_{ij}^{(1)} - \frac{\lambda_2}{\rho} w_{ij}^{(12)} - \frac{\lambda_2}{\rho} w_{ij}^{(13)} > A_{ij}^{(3)} - \frac{\lambda_2}{\rho} w_{ij}^{(23)} + \frac{\lambda_2}{\rho} w_{ij}^{(13)} > A_{ij}^{(2)} + \frac{\lambda_2}{\rho} w_{ij}^{(12)} + \frac{\lambda_2}{\rho} w_{ij}^{(23)}$, then

$$\left(\widehat{Z}_{ij}^{(1)}, \widehat{Z}_{ij}^{(2)}, \widehat{Z}_{ij}^{(3)}\right) = \left(A_{ij}^{(1)} - \frac{\lambda_2}{\rho} w_{ij}^{(12)} - \frac{\lambda_2}{\rho} w_{ij}^{(13)}, A_{ij}^{(2)} + \frac{\lambda_2}{\rho} w_{ij}^{(12)} + \frac{\lambda_2}{\rho} w_{ij}^{(23)}, A_{ij}^{(3)} - \frac{\lambda_2}{\rho} w_{ij}^{(23)} + \frac{\lambda_2}{\rho} w_{ij}^{(13)}\right);$$

(5) If $|A_{ij}^{(1)} - A_{ij}^{(3)} - \frac{\lambda_2}{\rho} w_{ij}^{(12)} + \frac{\lambda_2}{\rho} w_{ij}^{(23)}| \leq \frac{\lambda_2}{\rho} w_{ij}^{(13)}$, and

$$\frac{A_{ij}^{(1)} + A_{ij}^{(3)} - \frac{\lambda_2}{\rho}(w_{ij}^{(12)} + w_{ij}^{(23)})}{2} > A_{ij}^{(2)} + \frac{\lambda_2}{\rho}(w_{ij}^{(12)} + w_{ij}^{(23)}),$$

then

$$\begin{aligned} & \left(\widehat{Z}_{ij}^{(1)}, \widehat{Z}_{ij}^{(2)}, \widehat{Z}_{ij}^{(3)}\right) \\ = & \left(\frac{A_{ij}^{(1)} + A_{ij}^{(3)} - \frac{\lambda_2}{\rho}(w_{ij}^{(12)} + w_{ij}^{(23)})}{2}, A_{ij}^{(2)} + \frac{\lambda_2}{\rho}(w_{ij}^{(12)} + w_{ij}^{(23)}), \frac{A_{ij}^{(1)} + A_{ij}^{(3)} - \frac{\lambda_2}{\rho}(w_{ij}^{(12)} + w_{ij}^{(23)})}{2}\right); \end{aligned}$$

(6) If $A_{ij}^{(2)} - \frac{\lambda_2}{\rho} w_{ij}^{(12)} - \frac{\lambda_2}{\rho} w_{ij}^{(23)} > A_{ij}^{(1)} + \frac{\lambda_2}{\rho} w_{ij}^{(12)} - \frac{\lambda_2}{\rho} w_{ij}^{(13)} > A_{ij}^{(3)} + \frac{\lambda_2}{\rho} w_{ij}^{(23)} + \frac{\lambda_2}{\rho} w_{ij}^{(13)}$, then

$$\left(\widehat{Z}_{ij}^{(1)}, \widehat{Z}_{ij}^{(2)}, \widehat{Z}_{ij}^{(3)}\right) = \left(A_{ij}^{(1)} + \frac{\lambda_2}{\rho} w_{ij}^{(12)} - \frac{\lambda_2}{\rho} w_{ij}^{(13)}, A_{ij}^{(2)} - \frac{\lambda_2}{\rho} w_{ij}^{(12)} - \frac{\lambda_2}{\rho} w_{ij}^{(23)}, A_{ij}^{(3)} + \frac{\lambda_2}{\rho} w_{ij}^{(23)} + \frac{\lambda_2}{\rho} w_{ij}^{(13)}\right);$$

(7) If $|A_{ij}^{(1)} - A_{ij}^{(3)} + \frac{\lambda_2}{\rho} w_{ij}^{(12)} - \frac{\lambda_2}{\rho} w_{ij}^{(23)}| \leq \frac{\lambda_2}{\rho} w_{ij}^{(13)}$, and

$$\frac{A_{ij}^{(1)} + A_{ij}^{(3)} + \frac{\lambda_2}{\rho}(w_{ij}^{(12)} + w_{ij}^{(23)})}{2} < A_{ij}^{(2)} - \frac{\lambda_2}{\rho}(w_{ij}^{(12)} + w_{ij}^{(23)}),$$

then

$$
\begin{aligned}
&\left(\widehat{Z}_{ij}^{(1)}, \widehat{Z}_{ij}^{(2)}, \widehat{Z}_{ij}^{(3)}\right) \\
= &\left(\frac{A_{ij}^{(1)} + A_{ij}^{(3)} + \frac{\lambda_2}{\rho}(w_{ij}^{(12)} + w_{ij}^{(23)})}{2}, A_{ij}^{(2)} - \frac{\lambda_2}{\rho}(w_{ij}^{(12)} + w_{ij}^{(23)}), \frac{A_{ij}^{(1)} + A_{ij}^{(3)} + \frac{\lambda_2}{\rho}(w_{ij}^{(12)} + w_{ij}^{(23)})}{2}\right);
\end{aligned}
$$

(8) If $A_{ij}^{(2)} - \frac{\lambda_2}{\rho} w_{ij}^{(12)} - \frac{\lambda_2}{\rho} w_{ij}^{(23)} > A_{ij}^{(3)} + \frac{\lambda_2}{\rho} w_{ij}^{(23)} - \frac{\lambda_2}{\rho} w_{ij}^{(13)} > A_{ij}^{(1)} + \frac{\lambda_2}{\rho} w_{ij}^{(12)} + \frac{\lambda_2}{\rho} w_{ij}^{(13)}$, then

$$
\left(\widehat{Z}_{ij}^{(1)}, \widehat{Z}_{ij}^{(2)}, \widehat{Z}_{ij}^{(3)}\right) = \left(A_{ij}^{(1)} + \frac{\lambda_2}{\rho} w_{ij}^{(12)} + \frac{\lambda_2}{\rho} w_{ij}^{(13)}, A_{ij}^{(2)} - \frac{\lambda_2}{\rho} w_{ij}^{(12)} - \frac{\lambda_2}{\rho} w_{ij}^{(23)}, A_{ij}^{(3)} + \frac{\lambda_2}{\rho} w_{ij}^{(23)} - \frac{\lambda_2}{\rho} w_{ij}^{(13)}\right);
$$

(9) If $|A_{ij}^{(2)} - A_{ij}^{(3)} - \frac{\lambda_2}{\rho} w_{ij}^{(12)} + \frac{\lambda_2}{\rho} w_{ij}^{(13)}| \leq \frac{\lambda_2}{\rho} w_{ij}^{(23)}$, and

$$
\frac{A_{ij}^{(2)} + A_{ij}^{(3)} - \frac{\lambda_2}{\rho}(w_{ij}^{(12)} + w_{ij}^{(13)})}{2} > A_{ij}^{(1)} + \frac{\lambda_2}{\rho}(w_{ij}^{(12)} + w_{ij}^{(13)}),
$$

then

$$
\begin{aligned}
&\left(\widehat{Z}_{ij}^{(1)}, \widehat{Z}_{ij}^{(2)}, \widehat{Z}_{ij}^{(3)}\right) \\
= &\left(A_{ij}^{(1)} + \frac{\lambda_2}{\rho}(w_{ij}^{(12)} + w_{ij}^{(13)}), \frac{A_{ij}^{(2)} + A_{ij}^{(3)} - \frac{\lambda_2}{\rho}(w_{ij}^{(12)} + w_{ij}^{(13)})}{2}, \frac{A_{ij}^{(2)} + A_{ij}^{(3)} - \frac{\lambda_2}{\rho}(w_{ij}^{(12)} + w_{ij}^{(13)})}{2}\right);
\end{aligned}
$$

(10) If $A_{ij}^{(3)} - \frac{\lambda_2}{\rho} w_{ij}^{(23)} - \frac{\lambda_2}{\rho} w_{ij}^{(13)} > A_{ij}^{(1)} - \frac{\lambda_2}{\rho} w_{ij}^{(12)} + \frac{\lambda_2}{\rho} w_{ij}^{(13)} > A_{ij}^{(2)} + \frac{\lambda_2}{\rho} w_{ij}^{(12)} + \frac{\lambda_2}{\rho} w_{ij}^{(23)}$, then

$$
\left(\widehat{Z}_{ij}^{(1)}, \widehat{Z}_{ij}^{(2)}, \widehat{Z}_{ij}^{(3)}\right) = \left(A_{ij}^{(1)} - \frac{\lambda_2}{\rho} w_{ij}^{(12)} + \frac{\lambda_2}{\rho} w_{ij}^{(13)}, A_{ij}^{(2)} + \frac{\lambda_2}{\rho} w_{ij}^{(12)} + \frac{\lambda_2}{\rho} w_{ij}^{(23)}, A_{ij}^{(3)} - \frac{\lambda_2}{\rho} w_{ij}^{(23)} - \frac{\lambda_2}{\rho} w_{ij}^{(13)}\right);
$$

(11) If $|A_{ij}^{(1)} - A_{ij}^{(2)} + \frac{\lambda_2}{\rho} w_{ij}^{(13)} - \frac{\lambda_2}{\rho} w_{ij}^{(23)}| \leq \frac{\lambda_2}{\rho} w_{ij}^{(12)}$, and

$$
\frac{A_{ij}^{(1)} + A_{ij}^{(2)} + \frac{\lambda_2}{\rho}(w_{ij}^{(13)} + w_{ij}^{(23)})}{2} < A_{ij}^{(3)} - \frac{\lambda_2}{\rho}(w_{ij}^{(23)} - w_{ij}^{(13)}),
$$

then

$$
\begin{aligned}
&\left(\widehat{Z}_{ij}^{(1)}, \widehat{Z}_{ij}^{(2)}, \widehat{Z}_{ij}^{(3)}\right) \\
= &\left(\frac{A_{ij}^{(1)} + A_{ij}^{(2)} + \frac{\lambda_2}{\rho}(w_{ij}^{(13)} + w_{ij}^{(23)})}{2}, \frac{A_{ij}^{(1)} + A_{ij}^{(2)} + \frac{\lambda_2}{\rho}(w_{ij}^{(13)} + w_{ij}^{(23)})}{2}, A_{ij}^{(3)} - \frac{\lambda_2}{\rho}(w_{ij}^{(23)} - w_{ij}^{(13)})\right);
\end{aligned}
$$

(12) If $A_{ij}^{(3)} - \frac{\lambda_2}{\rho}w_{ij}^{(23)} - \frac{\lambda_2}{\rho}w_{ij}^{(13)} > A_{ij}^{(2)} - \frac{\lambda_2}{\rho}w_{ij}^{(12)} + \frac{\lambda_2}{\rho}w_{ij}^{(23)} > A_{ij}^{(1)} + \frac{\lambda_2}{\rho}w_{ij}^{(12)} + \frac{\lambda_2}{\rho}w_{ij}^{(13)}$, then

$$\left(\widehat{Z}_{ij}^{(1)}, \widehat{Z}_{ij}^{(2)}, \widehat{Z}_{ij}^{(3)}\right) = \left(A_{ij}^{(1)} + \frac{\lambda_2}{\rho}w_{ij}^{(12)} + \frac{\lambda_2}{\rho}w_{ij}^{(13)}, A_{ij}^{(2)} - \frac{\lambda_2}{\rho}w_{ij}^{(12)} + \frac{\lambda_2}{\rho}w_{ij}^{(23)}, A_{ij}^{(3)} - \frac{\lambda_2}{\rho}w_{ij}^{(23)} - \frac{\lambda_2}{\rho}w_{ij}^{(13)}\right);$$

(13) If

$$\begin{aligned} |A_{ij}^{(1)} + A_{ij}^{(2)} - 2A_{ij}^{(3)}| &\leq \frac{3\lambda_2}{\rho}(w_{ij}^{(13)} + w_{ij}^{(23)}), \\ |A_{ij}^{(1)} + A_{ij}^{(3)} - 2A_{ij}^{(2)}| &\leq \frac{3\lambda_2}{\rho}(w_{ij}^{(12)} + w_{ij}^{(23)}), \\ |A_{ij}^{(2)} + A_{ij}^{(3)} - 2A_{ij}^{(1)}| &\leq \frac{3\lambda_2}{\rho}(w_{ij}^{(12)} + w_{ij}^{(13)}), \end{aligned}$$

then

$$\left(\widehat{Z}_{ij}^{(1)}, \widehat{Z}_{ij}^{(2)}, \widehat{Z}_{ij}^{(3)}\right) = \left(\frac{A_{ij}^{(1)} + A_{ij}^{(2)} + A_{ij}^{(3)}}{3}, \frac{A_{ij}^{(1)} + A_{ij}^{(2)} + A_{ij}^{(3)}}{3}, \frac{A_{ij}^{(1)} + A_{ij}^{(2)} + A_{ij}^{(3)}}{3}\right).$$

### References


Boyd, S., Parikh, N., Chu, E., Peleato, B. and Eckstein, J. (2010). Distributed optimization and statistical learning via the alternating direction method of multipliers. *Foundns Trends Mach. Learn.*, 3, 1–122.

Danaher, P., Wang, P. and Witten, D. M. (2014). The joint graphical lasso for inverse covariance estimation across multiple classes. *J.R.Statist.Soc.B*, 76, 373–397

Friedman, J., Hastie, T., Hoefling, H. and Tibshirani, R. (2007). Pathwise coordinate optimization. *Ann. Appl. Statist.*, 1, 302–332.

Tibshirani, R., Saunders, M., Rosset, S., Zhu, J. and Knight, K. (2005). Sparsity and smoothness via the fused lasso. *J. R. Statist. Soc. B*, 67, 91–108.
